# Divergent biosynthesis of monoterpene indole alkaloids from geissoschizine

**DOI:** 10.1101/2024.10.22.619577

**Authors:** Mohamed O. Kamileen, Benke Hong, Klaus Gase, Maritta Kunert, Lorenzo Caputi, Benjamin R. Lichman, Sarah E. O’Connor

## Abstract

Plants can generate structural diversity by enzymatic rearrangement of a central intermediate. 19*E*-geisssochizine is one such chemically versatile intermediate that plays a central role in the biosynthesis of monoterpene indole alkaloids such as strychnine, ibogaine and vinblastine. Here we report how 19*E*-geissoschizine undergoes oxidative transformations to generate four distinct alkaloid scaffolds through the action of three biosynthetic enzymes. Using in vitro enzymatic assays and gene silencing, we demonstrate how these three cytochrome P450 enzymes in the medicinal plant *Catharanthus roseus* transform 19*E*-geisssochzine into *strychnos, sarpagan, akuammiline*-type, and *mavacurane-*type alkaloids. We use mutational analysis to show how minimal changes to the active site of these similar enzymes modulate product specificity. This work highlights how substrate reactivity and enzyme mutations work synergistically to generate chemical diversity.

---

The monoterpene indole alkaloids are a large family of ca. 3000 structurally diverse alkaloids originating from a single substrate, strictosidine^1^. Typically, strictosidine is deglucosylated by strictosidine glucosidase (SGD) and then subsequently reduced to form 19*E*-geissoschizine (geissoschizine) by the medium-chain alcohol dehydrogenase geissoschizine synthase (GS)^2–4^. Geissoschizine, a *corynanthe-type* alkaloid, is densely functionalized, harboring an alpha-beta unsaturated carbonyl, enol, indole and basic nitrogen, allowing a wide range of downstream reactions^5^ and is the precursor for *strychnos, sarpagan, akuammiline*, and *mavacurane*-type monoterpene indole alkaloids^5,6^ (**Figure 1**). Here, we show how three cytochrome P450 enzyme homologs oxidize geissoschizine to form these four distinct alkaloid scaffolds, and how the activity of these enzymes can be interconverted through mutations in the active site. The results demonstrate how nature capitalizes on the inherent reactivity of a highly versatile starting substrate to generate a variety of natural product scaffolds.

**Figure 1.**
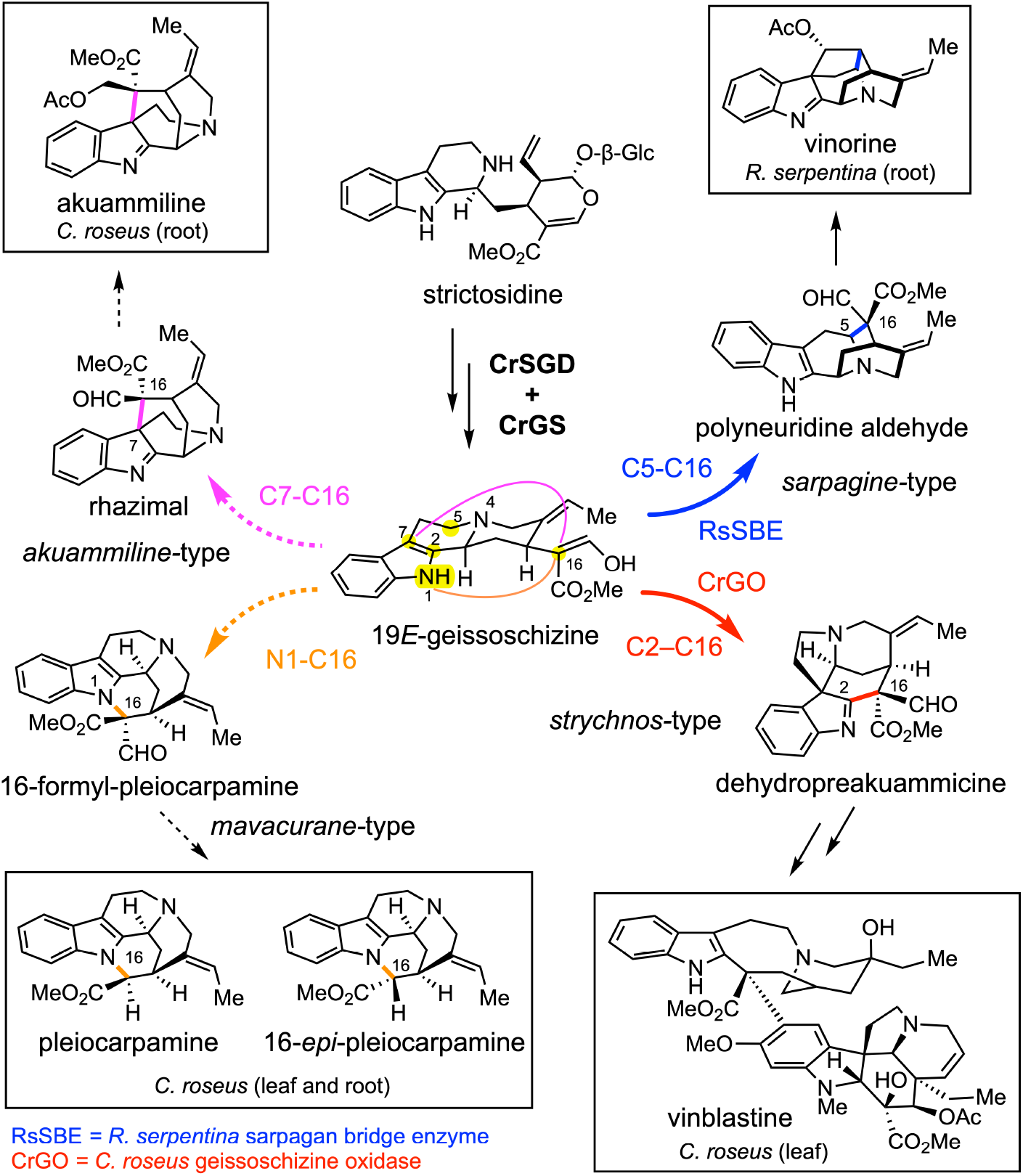
Proposed oxidative transformation of 19*E*-geissoschizine into four scaffolds.

Previously reported enzymes that oxidize geissoschizine belong to the cytochrome P450 CYP71 subfamily^2,7–10^. The C16-formyl ester moiety and the C16*R* stereocenter of geissoschizine play a crucial role in downstream scaffold biosynthesis^2,9,11^. The *strychnos*-type scaffold dehydropreakuammicine is formed by oxidative coupling of C16 and C2 by the *Catharanthus roseus* CYP71 enzyme CrGO (geissoschizine oxidase, also identified in *Strychnos nux vomica*)^2,9^, while SBE (sarpagan bridge enzyme), isolated from the related plant *Rauvolfia serpentina*, catalyzes formation of the C16 and C5 bond to form the *sarpagan* scaffold, polyneuridine aldehyde^7^ (**Figure 1**). RS (rhazimal synthase), which catalyzes formation of the C16 and C7 bond to form the *akuammiline*-type alkaloid rhazimal, has been identified in *Alstonia scholaris*^9^. In principle, the C16 carbon of geissoschizine can also react with the indole nitrogen to form the *mavacurane*-type alkaloids pleiocarpamine^5^, though an enzyme that catalyzes this transformation has not been identified (**Figure 1**). The medicinal plant *C. roseus* is only known for production of alkaloids derived from the GO product, dehydropreakuammicine (*strychnos*-type)^2^. However, we noted that the transcriptome of *C. roseus* encodes numerous GO homologs with unknown function.

We curated and built a phylogeny consisting of these 16 unknown *C. roseus* CYP71 genes along with similar P450s of known function (**Figure S1)**. The CYP71 enzymes of unknown function were expressed in *Saccharomyces cerevisiae* (yeast), and isolated microsomes containing the heterologous P450 were screened for activity with geissoschizine. Candidate CRO_T014436 yielded a major product (*m/z* 351.1703), a minor product with (*m/z* 323.1754) and trace amounts of akuammicine (*m/z* 323.1754), which is the deformylation product of dehydropreakuammicine (*strychnos-*type) (**Figure 2a**)^2^. To identify the major and minor products, we scaled up the in vitro reaction for isolation. The minor product was shown by NMR analysis to be the *akuammiline*-type alkaloid strictamine, which is the deformylation product of rhazimal (**Supporting NMR S1-S6**). The major product converted to strictamine over the course of purification, but partial NMR analysis of this unstable product was consistent with a structural assignment of rhazimal (**Supporting NMR S7-S9**) (**Figure 2a**). Therefore, we named this enzyme rhazimal synthase (CrRS). Rhazimal, strictamine or alkaloids derived from these products have not been reported from *C. roseus*. However, when we carefully profiled the roots and leaves of *C. roseus* in search of *akuammiline*-type alkaloids, we found the rhazimal derived compound, akuammiline, in the roots (**Figure S2**). This suggests that rhazimal in *C. roseus* is metabolized into akuammiline. In the absence of downstream enzymes, deformylation of rhazimal to strictamine occurs. The *in planta* function of CrRS could not be definitively established by virus-induced gene silencing (VIGS), since CrRS expressed in the roots and silencing is only possible in *C. roseus* leaf and stem tissue.

**Figure 2.**
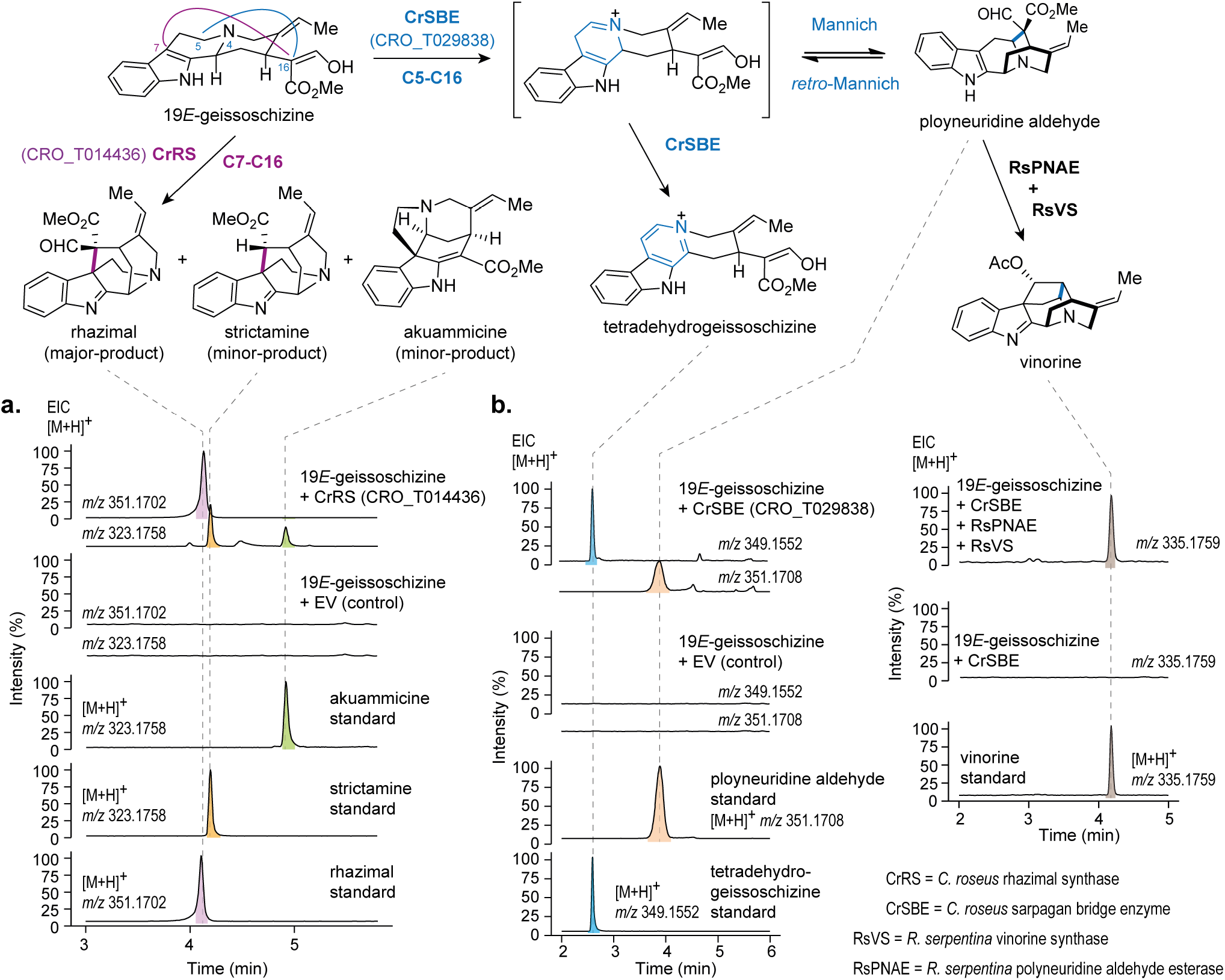
Oxidative transformations of geissoschizine in *C. roseus*. (**a**) Biosynthesis of rhazimal from CrRS. RS catalyzes an oxidation to forge the C7-C16 bond by oxidative coupling 19*E*-geissoschizine. Rhazimal readily deformylates to form strictamine. (**b**) Biosynthesis of polyneuridine aldehyde and tetradehydrogeissoschizine from CrSBE. SBE oxidizes 19*E*-geissoschizine to form the iminium ion at C5-N4, which then allows a Mannich reaction to form the C5-C16 bond to generate polyneuridine aldehyde. Alternatively, the initial iminium product oxidizes to form the aromatic product tetradehydrogeissoschizine. Polyneuridine aldehyde can be captured by downstream enzymes (RsPNAE, RsVS) that convert it to the stable product vinorine.

Assay of CRO_T029838 with geissoschizine led to production of an oxidized product that co-eluted with a synthetic tetradehydrogeissoschizine (*m/z* 349.1547) standard. We could also detect small amounts of polyneuridine aldehyde (*m/z* 351.1703), which also co-eluted with the respective standard (**Figure 2b**). We named this enzyme sarpagan bridge enzyme (CrSBE). Notably, polyneuridine aldehyde appeared to be the initially formed product, but was converted to tetradehydrogeissoschizine over time (**Figure S3**). Polyneuridine aldehyde synthesis most likely takes place via an initial oxidation of the C5-N4 bond to form an iminium species. Formation of polyneuridine aldehyde involves bond formation between C16 and C5 via a Mannich reaction. We hypothesized that this C5-C16 bond forming reaction can take place in reverse, reverting to the initial oxidation iminium ion product by a retro-Mannich reaction. Therefore, formation of polyneuridine aldehyde would be a reversible process. Formation of tetradehydrogeissoschizine is formed by a second oxidation of the initial iminium intermediate formed by SBE. This process, in contrast, is likely to be irreversible, due to the formation of the highly stable aromatic β-carboline moiety (**Figure 2b**). Therefore, the initially formed polyneuridine aldehyde product would gradually convert to tetradehydrogeissoschizine over time. To explore how the polyneuridine aldehyde product could be funneled into downstream metabolic processes, we assayed CrSBE in combination with previously characterized enzymes known to convert polyneuridine aldehyde to the stable compound vinorine by the enzymes polyneuridine aldehyde esterase (PNAE)^12^ and vinorine synthase (VS)^13,14^ from *R. serpentina*. The clean formation of vinorine (*m/z* 335.1752) (**Figure 2c**) demonstrates how the Mannich reaction is favored when downstream enzymes are present to consume the initially formed polyneuridine aldehyde. Notably, *C. roseus* produces no detectable products derived from polyneuridine aldehyde, dehydrogeissoschizine or tetradehydrogeissoschizine. We speculated that CrSBE may oxidize the *corynanthe* alkaloids ajmalicine/tetrahydroalstonine, substrates that are structurally similar to geissoschizine, *in planta*. However, when CrSBE was silenced in *C. roseus* leaves by VIGS, there was no significant reduction in the levels of the oxidized products (serpentine/alstonine) (**Figure S4**). The physiological function of CrSBE in *C. roseus* therefore remains unknown.

None of the remaining 14 CYP candidates showed activity with geissoschizine. Therefore, with three functional *C. roseus* CYPs in hand (CrGO, CrRS, CrSBE, 51-75 % amino acid identity, **Figure S5, S6**), we set out to understand the molecular basis of this product selectivity. Structural models of CrGO, CrRS and CrSBE were constructed and compared to identify amino acids that could be responsible for the cyclization specificity, which were then targeted for mutational analysis (**Figure 3a**). Using this approach, the native activity of CrRS (production of *akuammiline*-type alkaloids rhazimal/strictamine) could be converted to GO activity (production of the *strychnos*-type alkaloids dehydropreakuammicine/akuammicine) as demonstrated by the activity of mutants CrRS-M1 (4 mutations) and CrRS-M2 (10 mutations) (**Figure 3b**). The native activity of CrRS could also be converted to SBE activity (production of *sarpagan*-type alkaloid polyneuridine aldehyde along with tetradehydrogeissoschizine) with CrRS M3 (4 mutations) and CrRS-M4 (5 mutations) (**Figure 3c**). CrGO could be partially switched to CrRS (CrGO-M1) and to CrSBE (CrGO-M2) (**Figure S7**) while CrSBE could be partially switched to CrRS (CrSBE-M3 and M6) (**Figure S7**). Conversion of CrSBE to CrGO was not possible. This mutational analysis suggests that the active site of these enzymes can be reshaped with relatively few mutations, which in turns allows the flexible geissoschizine substrate to bind in alternative positions. Subsequent oxidation and cyclization would proceed with altered regioselectivity to generate the three different scaffolds.

**Figure 3.**
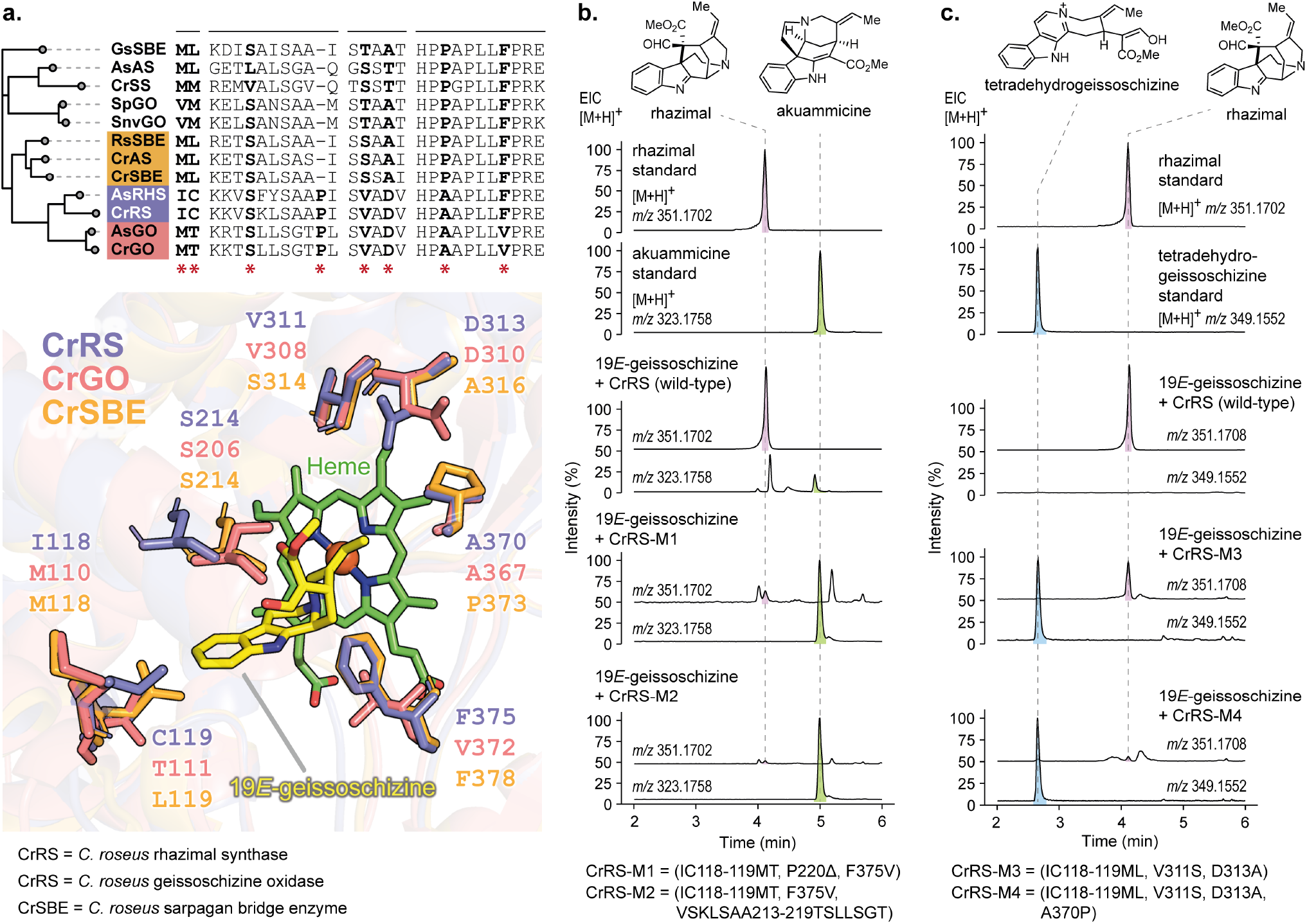
Site directed mutagenesis of geissoschizine oxidases. (**a**) Phylogeny and amino acid residues (stars) surrounding the catalytic pocket of SBE, GO, and RS enzymes. Residues selected for targeted mutagenesis in the active site. 19*E*-geissoschizine is shown in yellow. (**b**) Switch of CrRS to GO (mutants CrRS-M1 and M2). (**c**) Switch of CrRS to SBE (mutants CrRS-M3 and M4). The tetradehydrogeissoschizine product of CrSBE is observed under these assay conditions. Activity of additional mutants is shown the **Figure S7**.

The last missing predicted enzyme would catalyze cyclization via N1-C16 bond formation. This would result in formation of 16-formyl-pleocarpamine, which, like the RS product rhazimal and the GO product dehydropreakuammicine, is expected to undergo spontaneous deformylation, in this case producing pleiocarpamine and 16-*epi*-pleocarpamine^15^ (**Figure 4a**). However, none of the 16 CYP candidates or mutants appeared to produce 16-formyl-pleocarpamine, pleiocarpamine or 16-*epi*-pleocarpamine. Upon careful inspection, we could detect the presence of pleiocarpamine and 16-*epi*-pleocarpamine in *C. roseus* leaves and root, which strongly suggested that *C. roseus* should harbor an enzyme with this cyclization specificity (**Figure 4b, Figure S8**). To our surprise, when we closely examined the product profile of CrGO *in vitro* reaction assays, trace but detectable amounts of 16-*epi*-pleocarpamine along with the major, previously reported product akuammicine^2^ were observed (**Figure 4c**). This minor product shared identical MS^2^ and retention time to an authentic standard of 16-*epi*-pleocarpamine, which suggested deformylation from an initially formed 16-formyl-pleocarpamine product. The same results were observed when CrGO was expressed in leaves of *Nicotiana benthamiana*, and disks of these leaf tissues were incubated with geissoschizine (**Figure 4c**).

**Figure 4.**
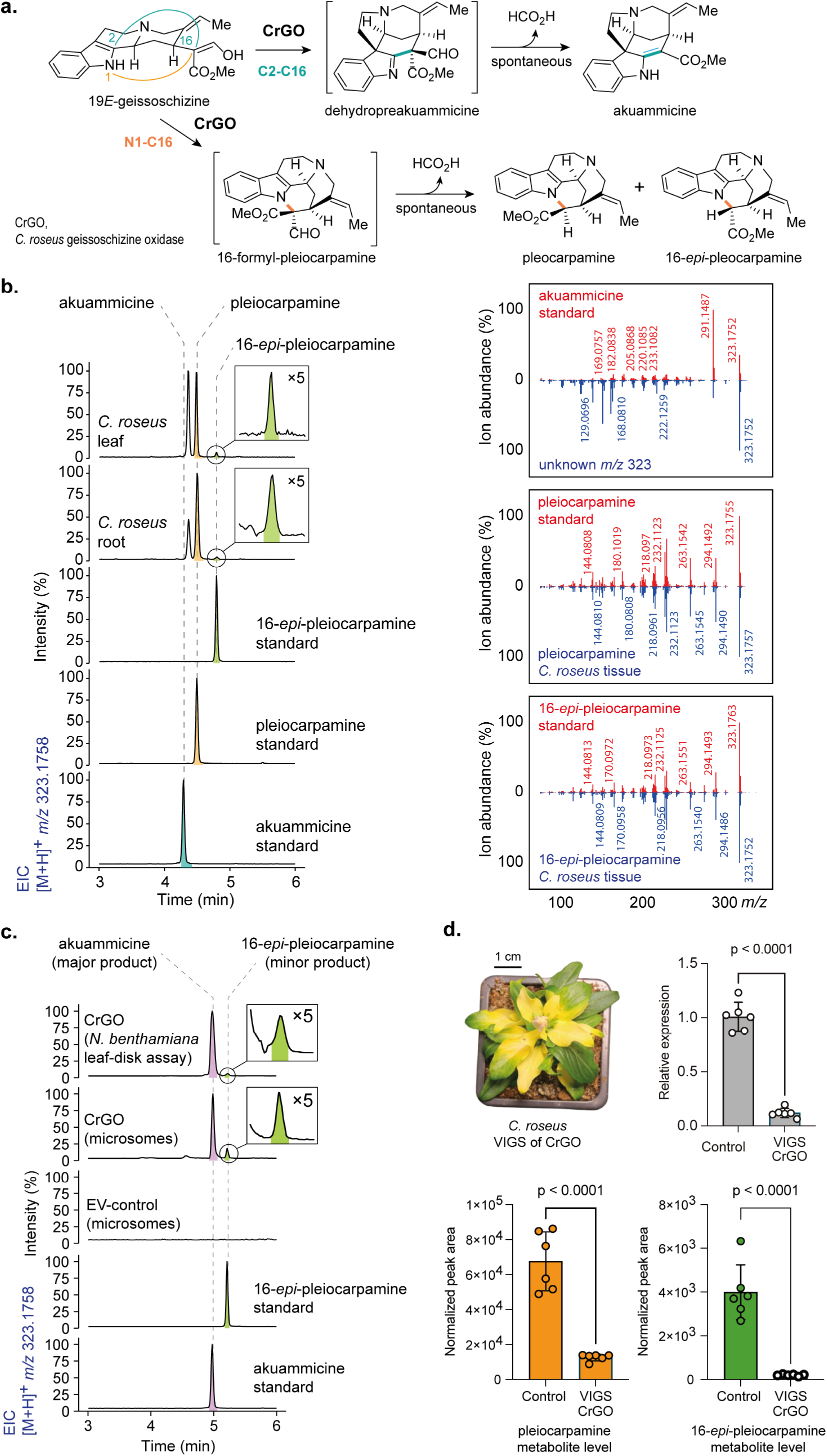
Biosynthesis of *mavacurane*-type alkaloids pleiocarpamine and 16-*epi*-pleiocarpamine in *C. roseus*. (**a**) Proposed biosynthesis of *mavacurane*-type and *strychnos*-type alkaloids. (**b**) Metabolite profiling of *mavacurane-type* alkaloids in *C. roseus* roots and leaf alongside the MS^2^ spectrum of authentic standards. (**c**) In vitro and in vivo activity of CrGO in producing 16-*epi*-pleiocarpamine. CrGO catalyzes oxidation of 19*E*-gesissoschizine to akuammicine forming C2-C16 bond. A minor product 16-*epi*-pleiocarpamine is observed forging the core mavacurane N1–C16 bond. (**d**) Silencing CrGO in *C. roseus* by VIGS leads to significant decrease of the *mavacurane*-type alkaloids. Bar graphs represents the values of the mean ± standard deviation (SD), p values present statistical analysis of two-tailed Student’s *t*-test. Extracted ion chromatograms (EIC) are presented with MS^2^ spectra displaying the fragmentation of the parent [M+H]^+^ ion.

We initially suspected that these low levels of 16-*epi*-pleocarpamine were simply an artifact of the *in vitro* enzymatic reaction. However, when we silenced CrGO in *C. roseus* using VIGS, a significant reduction in the levels of pleiocarpamine and 16-*epi*-pleocarpamine in *C. roseus* leaf was observed (**Figure 4d, S9**). Therefore, the minor product of this enzymatic reaction may play a physiologically significant role. Although it has been reported that 16-*epi*-pleocarpamine is the more thermostable epimer^11^, it is not clear why 16-*epi*-pleocarpamine predominates in the *in vitro* enzyme assay (**Figure 4c**) and why in the plant, the kinetic product pleocarpamine is predominantly observed (**Figure 4b**). We speculate that the conditions under which this cyclization takes place must impact the specificity of the subsequent deformylation.

Here we report the discovery of two cytochrome P450 enzymes, CrRS (rhazimal synthase) and CrSBE (sarpagan bridge enzyme), from *C. roseus*. Although enzymes with these activities had been identified from other plant species, *C. roseus* was not reported to have alkaloids derived from the products of these enzymes. Subsequent analysis revealed the presence of a CrRS-derived alkaloid, but the function of CrSBE remains unknown. The sequences of these three enzymes could be compared to demonstrate which residues are responsible for the regioselectivity of the oxidation and cyclization of geissoschizine. Additionally, we show that CrGO (geissoschizine oxidase), in addition to catalyzing C16-C2 bond formation, also catalyzes the C16-N1 bond formation required for the formation of the *mavacurane*-type alkaloids pleiocarpamine and 16-*epi*-pleiocarpamine (**Figure 1**). *Mavacurane*-type alkaloids have received limited attention^16,17^ but are known to converted into a range of complex bisindole alkaloids^5,18^. Silencing of GO in *C. roseus* strongly suggests that, although pleiocarpamine and 16-*epi*-pleiocarpamine are only formed as minor products by GO, this enzyme may contribute to mavacurane biosynthesis in the *C. roseus* plant. This raises the intriguing possibility that the production of minor side products in enzyme reactions can play a significant role in shaping the evolution of metabolic diversity.

## ASSOCIATED CONTENT

### Supporting Information

Supplemental information: Additional experimental details, assays, materials, and methods, and NMR spectra described in this study.

### Accession Codes

Genbank accession numbers: MH213134 (CrGO), pending (CrRS), pending (CrSBE)

### Notes

The authors declare no competing financial interests.

## ACKNOWLEDGMENT

Polyneuridine aldehyde was a generous gift from Dr. Laurent Evanno (Univ. Paris-Sud, CNRS, Université Paris-Saclay, France), 16-*epi*-pleiocarpamine was a generous gift from Dr. Guillaume Vincent (Univ. Paris-Sud, CNRS, Université Paris-Saclay, France) and pleiocarpamine was a generous gift from Prof. Dr. Hartmut Laatsch (Georg-August-Universität Göttingen, Germany). We would like to thank Sarah Heinicke for assistance with mass spectrometry. We also thank the members of the Max Planck Institute for Chemical Ecology Research Greenhouse for their care and provision of *Nicotiana benthamiana* and *Catharanthus roseus* plants.

## Supporting Information

