## Supplementary material for "Divergent biosynthesis of monoterpene indole alkaloids from geissoschizine": SI Data

**Table of Contents**


| | |
|---|---|
| Materials and Methods | Supporting information pages 02-07 |
| Supporting Tables | Supporting information pages 08-12 |
| Supporting Figures | Supporting information pages 13-22 |
| Supporting NMR | Supporting information pages 23-28 |
| Supporting References | Supporting information pages 29-31 |

### Materials and Methods

#### Chemicals and reagents

Geissoschizine was synthesised as previously described by Hong[1]. Polyneuridine aldehyde was a kind gift from Dr. Laurent Evanno (Univ. Paris-Sud, CNRS, Université Paris-Saclay, France)[2]. 16-*epi*-pleiocarpamine was a kind gift from Dr. Guillaume Vincent (Univ. Paris-Sud, CNRS, Université Paris-Saclay, France)[3]. Pleiocarpamine was a kind gift from Prof. Dr. Hartmut Laatsch (Georg-August-Universität Göttingen, Germany)[4]. Rhazimal was synthesized according to Zhang[5]. Tetradehydrogeissoschizine was synthesized as described by Turpin[6]. Akuammicine was biosynthesized as previously described by Tatsis[7]. Akuammiline was purchased from biosynth (https://www.biosynth.com).

#### Plant growth

*Catharanthus roseus* (var. “Sunstorm Apricot”, SA) was germinated in a standard soil mix in a climate chamber under a 16 h light/8 h dark photoperiod at 23-25°C, and 60% relative humidity. *C. roseus* var. SA plants, aged six months, were used to harvest young leaves for RNA isolation and cDNA synthesis for gene amplification. For the VIGS experiments, *C. roseus* var. SA 6-10 plants, 4 weeks old, were used. The *Nicotiana benthamiana* plants used for transient gene expression were grown in a greenhouse on a standard soil mix under a 16 h light/8 h dark photoperiod at 22 °C and 60% relative humidity. Plants were grown for 3-4 weeks before *Agrobacterium* infiltration. All plants were watered periodically before and after experiments.

#### Gene candidate selection

Publicly available *C. roseus* var. SA (Medicinal Plant Genomics Resource) was used to mine for cytochrome P450 based on homology to known CYP71AY genes involved in MIA biosynthesis. CYP71AY genes, AsAS (*Alstonia scholaris* alstonine synthase, OM323327.1)[8], AsGO (*Alstonia scholaris* geissoschizine oxidase, OM323328.1)[8], AsRHS *(Alstonia scholaris* rhazimal synthase, OM323329.1)[8], CrAS (*Catharanthus roseus* alstonine synthase, W8JDE2.1)[9], CrGO (*Catharanthus roseus* geissoschizine oxidase, JN613015.1)[7,10,11], CrSS (*Catharanthus roseus* serpentine synthase, MT829151.1)[12,13], GsSBE (*Gelsemium sempervirens* sarpagan bridge enzyme, MF537712)[9], RsSBE (*Rauvolfia serpentina* sarpagan bridge enzyme, MF537711)[9], SnvGO (*Strychnos nux-vomica* geissoschizine oxidase, OM304290.1)[1], SpGO (*Strychnos sp.* geissoschizine oxidase, OM304299.1)[1] were elected as bait to mine for uncharacterized CYP71AY genes from the *C. roseus* transcriptome. Full-length coding sequences (CDS) were curated, and MUSCLE[14] alignments were performed with *Arabidopsis thaliana* clan 86 genes as described in Kamileen[15]. A maximum-likelihood phylogenetic tree was created using IQ-TREE[16] under default parameters and visualized by iTOL[17]. Geneious Prime ver. 2023.1.2. was used for curation and analysis of transcriptomic data.

### Cloning methods

All genes described in this study (**Table S1**) were amplified by PCR from a cDNA library generated from *C. roseus* var. SA and *R. serpentina* using the primers listed in **Table S2**. The cDNA library for *C. roseus var.* SA was prepared by RNA isolation of 50 mg of young leaf using the RNeasy plant Mini kit (Qiagen) and converted to cDNA using SuperScript IV VILO reverse transcriptase (Thermo Fisher Scientific). P19-TBSV was cloned as previously described by Kamileen[18]. Site directed mutagenesis was performed using the overlap extension PCR strategy[19]. The codon(s) targeted for mutation were selected, and two overlapping primers for forward and reverse direction described in **Table S2** were designed to incorporate the desired mutation.

High-Fidelity Q5 Hot-Start 2X Master Mix (New England Biolabs) was used for gene amplification according to the manufacturer's instructions. Following PCR amplification, DNA products were analyzed on 1% agarose gel, and the excised gel fragments were purified using Zymoclean Gel DNA Recovery Kit (Zymo Research). In-Fusion HD cloning kit (Takara Bioscience) was used to ligate all amplified genes into target vectors according to the manufacturer's instructions.

For heterologous protein expression in yeast (*Saccharomyces cerevisiae,* WAT11), the purified PCR amplicons were cloned into pESC-HIS (GenBank: AF063850.1, Agilent) and chemically transformed into WAT11 cells as previously described by Kamileen[15]. For transient expression in *N. benthamiana*, the purified PCR amplicons were cloned into a modified 3Ω1[20] binary plasmid and transferred into *Agrobacterium tumefaciens* GV3101 cells by electroporation as previously described by Kamileen[15].

### Transient expression of candidate genes in *N. benthamiana*

The *Agrobacterium* strains transformed with the gene constructs of interest were grown overnight at 28°C with shaking at 220 rpm in volumes of 10 ml containing antibiotics 20 µg/ml rifampicin, 30 µg/ml gentamicin, 200 µg/ml spectinomycin. Cells were harvested by centrifugation at 4000 × *g* for 10 minutes, and the supernatant was discarded. The cell pellet was gently resuspended in infiltration buffer (10 mM MES, pH 5.6, 10 mM magnesium chloride, 200 µM acetosyringone). After a second centrifugation step, the supernatant was removed, and the pellet was resuspended in 10 ml of infiltration buffer, followed by incubation in the dark at room temperature with gentle rocking for 2 hours. Each *Agrobacterium* strain was adjusted to an OD600 of 0.3 and infiltrated into the abaxial side of a 3-week-old *N. benthamiana* leaf using a 1 ml needleless syringe. To account for batch variability, each experiment included at least three biological replicates, where individual leaves from three distinct plants were infiltrated. Wild-type and P19 controls were incorporated into the experimental design and tested both with and without substrate treatment. Leaves were harvested 4-days post-infiltration for subsequent plant disk assays.

### Heterologous expression of candidate genes in yeast

Yeast expression and microsomal preparation were performed as previously described by Kamileen[15]. Briefly, Yeast cells (WAT11) were cultured in 100 ml of SD-His medium containing 2% glucose (w/v) at 30°C, 200 rpm for 28–34 h. Cells were harvested and resuspended in 100 ml of SD-His medium + 1.8% galactose (w/v) + 0.2% glucose (w/v) for protein production at 30°C, 200 rpm for 18–24 h. Yeast cells were harvested, and microsomes were prepared as described by Pompon[21]. Prepared microsomes were stored at –80°C until assays were performed.

#### Enzyme assays with *Agrobacterium* infiltrated *N. benthamiana*

Enzyme assays using *N. benthamiana* leaf disk assays were performed as described by Kamileen[15]. Briefly, leaf disks were harvested from *N. benthamiana* three days post *Agrobacterium* infiltration. Each disk was incubated with 50 µM of substrate in a 250 µl volume of 50 mM HEPES buffer. The plates containing leaf disks were incubated in a growth chamber at 22 °C, 60% relative humidity and 16-h-light/8-h-dark photoperiod overnight (16-18h). Alternatively, 1 ml of 100 µM solution of geissoschizine substrate was infiltrated into the whole leaf 3 days post *Agrobacterium* infiltration. After 16-h incubation, the disks were harvested for metabolite extraction. A 250 µl extraction solution (70% methanol in water acidified with 0.1% formic acid) was added to the powdered tissue, mixed well by vortex and sonicated for 10 min at room temperature. The samples were then centrifuged at 20,000 × *g* for 10 min to pellet cell debris and the extract was diluted 1:2 with extraction solution. Samples were filtered through 0.22 µm PTFE syringe filters before metabolite analysis by LC-MS.

#### In vitro enzymatic assays

In vitro enzyme assays using yeast microsomal preparations containing heterologous protein were conducted in analytical-scale reactions, each with a total volume of 100 µl. The reactions were carried out in 50 mM HEPES buffer (pH 7.5) consisting of 50-60 µg of total microsomal protein (0.50 µg $\mu L^{-1}$ microsomal protein), 1 mM NADPH, and 25 µM substrate. Negative control reactions were performed with yeast microsomes from yeast that had been transformed with empty vector (EV). Reactions were initiated by the addition of substrate, gently vortexed, and incubating at 30 °C for 1 hour unless specified otherwise. The reactions were quenched with extraction buffer, centrifuged at 20 000 × *g* at room temperature for 10 min and filtered through 0.22-µm PTFE filters before LC-MS analysis.

#### Semi-preparative scale isolation of strictamine

Isolation of strictamine, the degradation product of rhazimal, was performed using CrRS yeast microsomes and 19*E*-geissoschzine substrate following the protocol described by Kamileen[15]. Briefly, yeast microsomes harboring CrRS were prepared from 1 L of culture using the conditions described above. A 50 ml reaction was set up to generate rhazimal and strictamine from 12 mg (0.034 mmol) of 19*E*-geissoschzine. The reaction consisted of 50 mM HEPES pH 7.5 buffer, 0.034 mmol NADPH. Reactions were initiated by the addition of 8 mg of yeast microsomes and incubated at incubated at 30°C, 200 rpm for 16 h. The reactions were terminated by the addition of 10 ml ethyl acetate. The aqueous reaction mixture was extracted three times with ethyl acetate and evaporated to dryness. Strictamine was then isolated by preparative TLC, as described previously by Zhang[5]. The resulting was then subjected to NMR analysis. Attempts to isolate and characterize enzymatically formed rhazimal were not successful, since rhazimal decomposed to strictamine during the isolation process, and therefore could not be fully characterized by NMR.

#### Virus induced gene silencing (VIGS) in *C. roseus*

The protocol for VIGS vector construction and VIGS generally followed the procedure described in Li[22]. In brief, 300 bp fragments (**Table S3**) of the coding regions of CrRS, CrSBE, and CrGO from the genomic DNA (gDNA) from *C. roseus var.* SA, by PCR amplification with primers designed for In-Fusion HD cloning (Takara bioscience) (**Table S2**) using Phusion™ High-Fidelity DNA Polymerase (Thermo Fischer scientific). Genomic DNA was isolated from *C. roseus* leaves with the DNeasy Plant Mini Kit (Qiagen). To avoid off-target gene silencing, the target regions for VIGS were selected using the SGN VIGS tool[23]. The oligonucleotide sequences are given in supplemental material. The obtained PCR fragments were cloned in the BamHI and, XhoI digested VIGS vector pTRV2-MgChl [pTRV2-ChlH(F1R2)][24] with the In-Fusion Snap Assembly Master Mix (Takara bioscience), yielding pTRV2-MgChl-CrGO and pTRV2-MgChl-CrSBE constructs.

*Agrobacterium tumefaciens* GV3101 was transformed with the three constructed plasmids and used to inoculate *C. roseus* var. SA. For this, *A. tumefaciens* GV3101 with plasmid pTRV1[25] and *A. tumefaciens* GV3101 carrying pTRV2-MgChl-CrGO, pTRV2-MgChl-CrSBE, and pTRV2-MgChl-CrRS were grown overnight in a rotary shaker at 28 °C and 300 rpm in each 10 ml of LB medium supplemented with 50 µg $ml^{-1}$ kanamycin, 25 µg $ml^{-1}$ gentamicin and 100 µg $ml^{-1}$ rifampicin to an OD600 of 2. Cells were pelleted for 10 min at 4000 × *g* and resuspended in infiltration buffer (100 µM acetosyringone, 10 mM NaCl and 1.75 mM $CaCl_2$) to an OD600 of 2. After 2 h of incubation on a rotary shaker at room temperature and 60 rpm, 450 µl of each bacterial strain containing a pTRV2 plasmid were mixed with the same volume of the bacterial strain harboring pTRV1. VIGS inoculation was performed by pipetting 10 µl of the mixed bacterial suspension between plant stem and petiole of one leaf first grown after the cotyledons of a 30 days old *C. roseus* var. SA plant. The stem was pierced with a ø 0.40 x 25 mm Sterican® needle twice through the bacterial suspension drop. Per pTRV2 construct 8 plants were inoculated; the plants inoculated with pTRV2-MgChl strain served as negative controls. After VIGS inoculation, plants were grown under *C. roseus* standard greenhouse conditions. First VIGS symptoms (yellowing of the leaves due to magnesium chelatase subunit H gene silencing) could be observed after 12 days. The leaf tissue with complete target gene silencing indicated by yellow colour was harvested three weeks after inoculation.

Leaves with a VIGS phenotype (bleaching) were carefully harvested from each experimental condition and the control plants from 6 independent biological replicates. The tissue was snap-frozen in liquid nitrogen and was ground to fine powder using tungsten beads. Total RNA extraction was performed using the RNeasy mini kit (Qiagen) according to the manufacturer’s instructions. The genomic DNA was digested, and the RNA was further purified using RNA Clean & Concentrator-25 with DNAse I (Zymo research). RNA (1000 ng) was reverse-transcribed using SuperScript-IV VILO master mix (Thermo Fisher Scientific) following manufacturer’s instructions. Gene-specific primers were designed to target 150-200 bp target of the gene of interest with the Primer BLAST software (NCBI). Primers designed for qPCR are listed in **Table S4**. The qPCR experiment was performed as previously described by Payne[26] using the delta delta cycle method using the 40S ribosomal protein 9 (Rps9) as a reference gene for normalization. The primer efficiency for the designed qPCR primers was between 95–102%. Measurements for qPCR analysis were performed using a QuantStudio 1 cycler (Applied Biosystems). Statistical analysis (two-tailed Student's *t*-test) was performed using Prism 10 (Version 10.2.2).

#### Metabolite analysis of *C. roseus* tissue

Nearly identical plants of *C. roseus* var. SA, 6 months of age, were selected for tissue profiling. The young leaves and roots of the plants were harvested and snap-frozen in liquid nitrogen. Tissue was pulverized to fine powder using a mortar and pestle, and 50 mg of tissue was extracted with 1 ml of the extraction solution (70% methanol in water acidified with 0.1% formic acid). Samples were then prepared for LC-MS analysis as described in the section, In vitro assays with *Agrobacterium* infiltrated *N. benthamiana* leaf disks.

#### LC-MS data acquisition and analysis of metabolites

LC-MS analysis was performed as described previously by Kamileen[18]. Samples were analyzed using a Thermo Scientific UltiMate 3000 ultra-high performance liquid chromatography (UHPLC) system coupled to an Impact II UHR-Q-ToF (Ultra-High Resolution Quadrupole-Time-of-Flight) mass spectrometer (Bruker Daltonics). Metabolites were separated by reversed-phase liquid chromatography using a Phenomenex Kinetex XB-C18 (100 x 2.1mm, 2.6 µm; 100 Å) column at 40 °C. The mobile phases for metabolite separation were water with 0.1% formic acid (A) and acetonitrile (B). A flow rate of 0.6 ml/min was used for the chromatography. The injection volume was 2 µl. The chromatographic separation was performed starting at 10% B for 1 min, linear gradient from 10% to 30% B in 6 min, 90% B for 1.5 min, 10% B for 2.5 min. Authentic standards were prepared as 20 µM solutions in methanol, and 2 µl was injected under the chromatographic conditions described above. Mass spectrometry acquisition was performed in positive electrospray ionization mode with a capillary voltage of 3500 V and an end plate offset of 500 V; a nebulizer pressure of 2.5 bar was used, with nitrogen at 250 °C and a flow of 11 l/min as the drying gas. Acquisition was done at 12 Hz in the mass range from 80 to 1000 *m/z*, with data dependent $MS^2$ and an active exclusion window of 0.2 min. For tandem mass spectroscopy ($MS^2$) fragmentation was triggered on an absolute threshold of 400 and limited to a total cycle time range of 0.5 s. For collision energy, the stepping option model (from 20 to 50 eV) was used. At the beginning of each sample run, a sodium formate-isopropanol calibration solution was directly infused to the source at 0.18 ml/hour using a syringe pump to calibrate MS spectra recorded. To avoid injection peak and salt contamination of the MS, the initial 1 min of the active chromatographic gradient of each run was discarded to waste.

For the analysis of 19*E*-geissoschizine, the LC method was modified as described by Hong[1]. The mobile phase (A) was changed to ammonia at 0.025% (pH 9.0), and the mobile phase (B) remained acetonitrile. The flow rate was set to 0.6 ml/min, and the gradient linearly increased from 15% to 50% acetonitrile in 4 minutes (from minutes 1 to 5), and the column was washed and equilibrated within 10 min. A Waters Acquity UPLC BEH-C18 column (2.1 x 50 mm; 1.7 µm) was used and was temperature controlled at 40°C). 2 µl of samples were injected. The MS conditions were not modified.

Data was visualized and analyzed using Bruker Compass Data Analysis (Version 5.3) software. For relative quantification and metabolomics, data was analyzed using default pipelines embedded in Bruker Compass MetaboScape 2021b software (version 7.0.1). The non-targeted metabolomics workflow was used to generate an output containing a list of mass signatures with retention times, along with qualitative peak intensities by automated integration of extracted ion chromatograms with 5 ppm tolerance, molecular formula determined by accurate mass of precursors and fragments and statistical test outputs (*P* value and two-tailed Student's *t*-tests).

**NMR analysis**

NMR spectra were measured on a 700 MHz Bruker Advance III HD spectrometer (Bruker Biospin GmbH, Rheinstetten, Germany). For spectrometer control and data processing Bruker TopSpin (version 3.6.1) was used. Deuterated chloroform ($CDCl_3$) was used as a solvent, and all NMR spectra were referenced to the residual solvent signals at $\delta_H$ 7.26 and $\delta_C$ 77.0.

**Homology modelling and docking**

Protein homology models were generated using both AlphaFold3[27] and SWISS-MODEL (https://swissmodel.expasy.org/)[28]. The catalytic heme was docked using AlphaFold3. The substrate 19*E*-geissoschizine was docked into the active site using Webina (AutoDock Vina, https://durrantlab.pitt.edu/webina/)[29]. Protein homology models were visualized and interpreted using PyMol (Version 2.5.5, Schrödinger, LLC).

**Supplementary Tables**

**Table S1.** Nucleotide sequences for genes cloned and described in this study.

| **Gene Name (GenBank Accession))** | **Nucleotide sequence** |
|---|---|
| CrGO<br>(JN613015.1) | **ATG**GAGTTTTCTTTCTCCTCACCAGCTCTCTACATAGTTTATTTCTTGCTTTTCTTTGTG<br>GTAAGGCAATTATTGAAACCCAAAAGTAAGAAAAAATTGCCACCAGGTCCAAGAACAC<br>TACCCTTAATTGGAAACCTTCATCAACTCTCGGGACCTTTACCTCATCGTACCCTAAAA<br>AATTTGTCCGATAAACATGGTCCTTTGATGCACGTGAAAATGGGCGAACGTTCGGCAA<br>TTATAGTATCAGATGCAAGAATGGCAAAAATAGTTCTTCATAATAACGGTTTAGCCGTT<br>GCAGATCGGTCAGTAAATACTGTCGCAAGTATTATGACTTATAATAGTTTGGGTGTTAC<br>CTTTGCTCAGTATGGAGATTACTTAACAAAATTACGTCAAATCTATACTTTGGAACTTTT<br>AAGTCAGAAAAAAGTTCGATCTTTCTACAGTTGTTTTGAAGATGAACTCGATACTTTTG<br>TTAAGTCAATTAAGTCTAACGTTGGACAACCTATGGTTTTGTACGAGAAAGCTTCTGCT<br>TATTTGTATGCTACCATTTGTAGAACTATTTTTGGGAGTGTTTGCAAGGAAAAGGAGAA<br>AATGATTAAGATAGTGAAGAAAACGTCGTTACTGTCTGGAACACCACTTAGATTGGAG<br>GATCTTTTTCCTTCTATGAGCATTTTTTGTAGATTTAGTAAAACTTTGAATCAATTGAGA<br>GGACTTTTGCAAGAAATGGATGATATTTTGGAAGAAATCATAGTTGAAAGGGAAAAGG<br>CTTCTGAAGTCTCAAAAGAGGCAAAGGATGATGAAGATATGCTTAGTGTTCTGTTGAG<br>GCATAAATGGTACAATCCAAGTGGTGCTAAATTCAGAATCACTAATGCAGACATCAAA<br>GCCATTATCTTTGAGTTGATCTTAGCAGCAACCCTAAGTGTAGCAGATGTTACAGAAT<br>GGGCTATGGTTGAAATTCTAAGGGATCCCAAAAGTTTGAAAAAAGTGTACGAAGAAGT<br>ACGAGGTATTTGTAAAGAAAAGAAAAGAGTGACAGGATACGACGTTGAAAAAATGGAA<br>TTTATGCGTCTTTGTGTGAAGGAATCAACAAGAATACATCCAGCTGCTCCTCTTTTAGT<br>TCCTAGAGAATGCAGAGAAGATTTTGAAGTAGATGGATACACAGTTCCAAAAGGGGCT<br>TGGGTTATAACAAATTGCTGGGCTGTTCAAATGGATCCTACAGTATGGCCCGAACCAG<br>AGAAGTTTGATCCAGAAAGATATATTCGTAACCCAATGGATTTCTATGGAAGTAACTTT<br>GAATTGATACCATTTGGGACAGGTAGAAGAGGATGTCCTGGTATTTTATATGGAGTGA<br>CAAATGCTGAGTTTATGTTGGCTGCTATGTTTTATCATTTTGATTGGGAAATAGCAGAT<br>GGTAAAAAACCAGAAGAAATTGATTTGACTGAGGACTTTGGTGCTGGTTGCATTATGA<br>AGTATCCTTTGAAATTGGTTCCTCATCTTGTTAACGATTAG |
| CrRS<br>(Genbank ID pending) | **ATG**GAAACTATCAACTTCTTCTCCTTCTCCTTTCCACCAGTGGTCTACTTTCTCACCTT<br>CATTGCCTTCCTTTTAGTAGCAAAAAGACTAATAAGTCCCAAAACCCACAAAAAATTAT<br>TACCACCAGGTCCATGGACCTTACCCTTCATTGGAAATTTACATCAGCTCCTAGCCGG<br>ATCATTACCACATAGAATCCTCAAAAATCTGTCCGATAAACACGGACCGCTGATGACA<br>ATAATGATGGGTGAAAGAACAACAATAATAGTGTCCTCCGCAAGAATGGCAAAATTGG<br>TGCTTCACACTAATGGTCTAGCAGTTGCCAATAGACCAATTAATAGTGTGGCAGAAAT<br>TATTTGTTACAATAATTTAGGCACAACTTTTGCAAAATATGGTGAATATTTAAAACAATT<br>ACGCCAACTTTACATGTCGGAACTTTTGAGCCCCAAAAGAGTAAAATCATTTTCGGGC<br>ATTTTTGAAGATGAACTTGAGAAATTCGTTGGTTCAATTAGATCTCAAGTTGGTAAAGA<br>AATGGTAATTTATTGGAAATCAACTGAATATTTATATTCAGCAATTTGTAGGGTCATGTT<br>TGGAAGTGTTTGTAGTGAAAGGGAAAAATTAATTAAAGTGTGTAAAAAAGTGTCGAAAT<br>TATCAGCAGCACCAATCAGGTTTGAAGATCTTTTTCCATCAATTAAAGGGCTTTCATTT<br>TTAACAGGAAGAGATAATGTTTTAAAAGGTTTGGTTAAAGATCTTGATGATGTTCTTGA<br>TATATTAATAGCTCAGCGTGAAAATGGTACTACTTTTGATGAAGGAGATATGCTTGGAC<br>TTCTATTGAAGATTAAGAGTGGTGAAATCTATAATGCCAAACTCCAGATCACAAATGAT<br>GATATCAAAGCCATCATTTTCGAGCTGATGTTAGCTGCAACTCTAAGCGTAGCAGACG<br>TAGTAGAATGGGCCATGGTTGAAATAATAAGATATCCACAAGTTTTGAAACAGGTGAA<br>AGAAGAATTGAATAAAGTATTAAATGGAAAACAAGAAATAAGAGGATCTGATCTAGAG<br>AAAATGGAATATCTTCATATGTGTGTTAAAGAATCGACCCGACTCCACCCAGCTGCCC<br>CTCTTCTATTCCCAAGAGAAGCCCGTGAAGAATTTCAAATTGATAACTACACAATTCCA<br>AAAGGTTCATGGATCATGACTAATTATTGGGCAATTGGGCGTGACCCTGAAATTTGGC<br>CTAAACCCGATGAATTTGATCCGGAAAGATTCAGAAATAGTAATATTGATTTTTATGGA<br>AACCATTTTGAATTAATTCCATTTGGTAGTGGAAGGAGAGGATGTCCAGGGATATTATT<br>TGGAATTACTGAAGCTGAATTTATATTGGCTGCTTTATTTTATCATTTTGATTGGAAACT<br>ACCAGGTGGAATGGACCCGGATCAGATTGATATGACTGAAGTATTCGGTGCGGGATG<br>TATTTTGAAAAATCCATTGACTGTGATACCGGTTGTTGCACAAGATTGA |

| | |
|---|---|
| CrSBE<br>(Genbank ID<br>pending) | **ATG**GATGAGATGATGAACTTCTCCCTCACCTCTCCCATTTCCTTTCTCCTCTCCTTACT<br>TCTTATCCTATTTCTTATCCTATTTATGATGAAAGGAAACCAAACCCACAATGGCAAAA<br>AATTACCACCGGGTCCCAAAAAACTTCCTATTATAGGAAACCTCCATCAAATGATAGG<br>TTCACATCCTCATCGTGTTCTAAAAAAATTGGCCGATAAATATGGACCTATTATGCACT<br>TGCAAATGGGCCAACGATCAGCCGTTGTTATATCATCAGCGGAAAAAGCAAAAGAAAT<br>ATTGAACATTTATGGTGTTCAAGTTGCTGATAGACCTCAACTCATCGTATCTAAAATTA<br>TGTTGTACAATAGTTTGGGTGTCACTTTTGCTCCTTATGGTGACTACTTAAAGCAATTG<br>CGTCAAATTTATGCCGTAGAACTTCTTAGCCCTAAAACAGTTAGGTCCTTTTGGACAAT<br>TATGGAAGATGAAGTTTCCACAATGGTTAACTCAATTAAATCTGAAATTGGTCAACCAA<br>TAATTTTGCATGATAAAATGATGACTTATTTGTATACTATGCTTTGTAGAGTTACATTTG<br>GTGGCGTATGTAATGGACGCGAGACACTAATAATGGCAGCTAAAGAAACGTCAGCGC<br>TTTCTGCTGCTATTAGGATTGAGGATTTGTTTCCTTCAGTGAAAATACTTCCTTTAATTA<br>GTGGATTAAAGTCAAGATTAAACAATTTGTTGAAAACACTTGATACCATTCTTGAGGAT<br>ATTATCAGTGTTCGTGAGAAGAAATTATTAACTCAGCCATTGTTGGATGATGAAGATAT<br>GTTGGGAGTTCTCCTCAAGTACAAAAATGGAAAGGGAAAAGATACTAAATTTAGAGTC<br>ACCAACAACGACATCAAGGCAGTCATTTTTGAAATTATCTTAGCTGGCACCCTAAGTT<br>CATCAGCTATAGTAGAATGGTGTATGTCTGAAATGATAAAAAACTCTGAAAAGTTTAAA<br>AAGGCACAAGATGAAGTTAGGAAAGTTTTAAAGGGTAAAAAAACAATTAGTGGAAGTG<br>ATGTTGGTAAAATGGAATATGTTAAAATGGTGGTTAAGGAATCTTTGAGATTACACCCT<br>CCTGCTCCCATATTGTTTCCAAGAGAATGTAGGGAAGAATTTGAGATAGATGGAATGA<br>CTATACCTAAAAAAACTTGGTTGATTGTAAACTACTGGCCAATAGGAAGAGATCCCAG<br>ACTTTGGGACGACGCCGACAAGTTTGAGCCGGAGAGGTTCAGTAATAGCAGCATTGA<br>TTTCAATGGAAGCCATTTTGAGTTGATACCATTTGGTGCTGGAAGAAGGATTTGTCCT<br>GGAATATTACTTGGAACAACAAATGTTGAGCTTTTACTTGCTACATTTCTCTATTATTTT<br>GATTGGAAACTTCCTCAAGGTATGAAACCCGAAGAATTAGATATGAATGAGGTATTTG<br>GTGCTGGTTGCATAAGGGAAAGTCCATTGTGTCTCATTCCTAGCATCTCATCAGCAGT<br>TGAAGGAAATTAA |
| RsPNAE<br>(AF178576.1) | **ATG**CATTCTGCTGCAAACGCCAAGCAACAAAAGCATTTTGTTCTGGTACACGGCGGAT<br>GTCTCGGAGCTTGGATCTGGTACAAGCTCAAGCCGCTGCTCGAGTCAGCCGGACATA<br>AGGTCACCGCCGTTGACCTGTCGGCCGCCGGCATCAACCCAAGAAGGCTCGATGAG<br>ATTCACACATTTCGGGACTACTCGGAGCCCTTGATGGAAGTCATGGCTAGTATTCCTC<br>CTGATGAGAAGGTTGTTCTTCTTGGCCATAGCTTTGGTGGCATGAGTTTGGGTCTTGC<br>CATGGAAACCTACCCAGAGAAGATATCAGTTGCTGTTTTTATGTCTGCAATGATGCCT<br>GATCCTAACCACTCACTAACTTATCCGTTTGAGAAGTACAATGAGAAGTGTCCGGCAG<br>ATATGATGTTGGACTCACAGTTTTCAACCTACGGAAACCCAGAGAACCCAGGAATGTC<br>AATGATTCTTGGACCTCAGTTTATGGCCCTCAAAATGTTCCAGAATTGCTCAGTCGAG<br>GACCTTGAATTAGCCAAAATGTTGACTCGACCAGGTTCGTTATTTTTCCAAGATTTGGC<br>CAAGGCCAAAAAGTTCTCAACCGAGAGGTACGGTTCGGTGAAGCGAGCTTATATCTTT<br>TGCAATGAAGATAAATCATTTCCAGTTGAGTTTCAGAAATGGTTTGTTGAAAGTGTTGG<br>AGCTGATAAAGTAAAAGAAATCAAAGAAGCAGATCATATGGGAATGCTTTCGCAGCCA<br>AGGGAAGTTTGCAAGTGCCTGCTTGATATATCAGATTCATAA |
| RsVS<br>(AJ556780.2) | **ATG**GCACCCCAGATGGAGAAAGTATCGGAGGAGCTGATTCTACCATCATCTCCAACA<br>CCCCAAAGCTTGAAATGCTATAAAATTTCCCACCTAGATCAACTGTTATTAACGTGTCA<br>CATCCCTTTTATTCTCTTCTATCCAAATCCGTTAGACTCAAACCTCGATCCTGCCCAGA<br>CATCTCAGCACCTGAAACAATCTTTGTCCAAAGTGTTAACTCACTTTTACCCTCTAGCT<br>GGAAGGATCAACGTAAATTCTTCCGTAGACTGTAATGATTCTGGAGTTCCTTTTGTCG<br>AAGCTCGGGTTCAAGCTCAACTCTCACAGGCAATTCAGAACGTCGTCGAGTTAGAAAA<br>ACTCGATCAATACCTTCCGTCCGCAGCTTATCCCGGCGGGAAAATTGAGGTGAACGA<br>GGATGTTCCCCTGGCTGTCAAAATCAGTTTCTTTGAGTGTGGAGGCACGGCCATTGG<br>TGTCAACTTATCGCATAAGATAGCTGATGTATTGTCCCTGGCCACCTTCCTCAACGCA<br>TGGACTGCCACATGCCGTGGGGAAACGGAGATTGTGCTACCTAATTTTGACTTGGCA<br>GCACGTCATTTTCCGCCCGTGGACAACACCCCGTCTCCTGAATTGGTACCGGATGAA<br>AACGTTGTGATGAAAAGATTCGTATTTGATAAAGAAAAAATAGGAGCCCTCAGAGCAC<br>AAGCTTCCTCGGCCTCAGAGGAGAAGAATTTCAGTCGGGTACAGCTTGTTGTTGCTTA<br>TATATGGAAGCACGTCATTGACGTGACCCGGGCAAAATATGGTGCTAAAAACAAGTTT<br>GTGGTAGTTCAAGCAGTGAACCTGAGGTCAAGAATGAATCCGCCCCTTCCTCACTAT<br>GCTATGGGGAACATCGCCACACTATTATTCGCGGCTGTAGATGCAGAGTGGGACAAA<br>GATTTTCCGGATCTCATCGGTCCGTTGAGAACCAGCCTAGAAAAAACTGAGGACGAC<br>CATAACCACGAATTACTAAAGGGAATGACTTGTTTGTATGAACTGGAACCTCAAGAAC<br>TTTTGTCTTTCACCAGTTGGTGTAGGCTTGGCTTTTATGACTTGGATTTCGGCTGGGG<br>GAAGCCTCTTTCAGCGTGCACAACAACTTTTCCCAAGAGGAACGCGGCGCTTTTGAT<br>GGATACAAGATCCGGAGATGGAGTGGAAGCATGGCTCCCAATGGCAGAAGATGAAAT<br>GGCGATGCTTCCTGTTGAATTGCTGTCACTTGTAGACAGCGATTTTAGCAAGTGA |

Notes: Start codon is highlighted in **bold**. Stop codon is underlined.

**Table S2.** Primers used in this study. Primer sequences. Primer sequences with homology to the destination plasmid for DNA assembly are in **bold**.

| Gene | Plasmid | Primer direction | Sequence (5‘-3‘) |
|---|---|---|---|
| *Nicotiana benthamiana* transient expression | | | |
| CrGO | 3Ω1 | Forward | **TTTATGAATTTTGCAGCTCG**ATGGAGTTTTCTTTCTCCTCA |
| | | Reverse | **GACAACCACAACAAGCACCG**CTAATCGTTAACAAGATGAGGAA |
| CrSBE | 3Ω1 | Forward | **TTTATGAATTTTGCAGCTCG**ATGGATGAGATGATGAACTTCTC |
| | | Reverse | **GACAACCACAACAAGCACCG**TTAATTTCCTTCAACTGCTGATGA |
| CrRS | 3Ω1 | Forward | **TTTATGAATTTTGCAGCTCG**ATGGAAACTATCAACTTCTTCTC |
| | | Reverse | **GACAACCACAACAAGCACCG**TCAATCTTGTGCAACAACCG |
| RsPNAE | 3Ω1 | Forward | **TTTATGAATTTTGCAGCTCG**ATGCATTCTGCTGCAAAC |
| | | Reverse | **GACAACCACAACAAGCACCG**TTATGAATCTGATATATCAAGCAGGC |
| RsVS | 3Ω1 | Forward | **TTTATGAATTTTGCAGCTCG**ATGGCACCCCAGATG |
| | | Reverse | **GACAACCACAACAAGCACCG**TCACTTGCTAAAATCGCTGT |
| p19-TBSV | 3Ω1 | Forward | **TTTATGAATTTTGCAGCTCG**ATGGAACGAGCTATACAAGGAAA |
| | | Reverse | **GACAACCACAACAAGCACCG**TTACTCGCTTTCTTCTTCGAAGG |
| **Gene** | **Plasmid** | **Primer direction** | **Sequence (5‘-3‘)** |
| *Saccharomyces cerevisiae* (WAT11) expression | | | |
| CrGO | pESC-HIS | Forward | **GAGAAAAAACCCCGGATCC**ATGGAGTTTTCTTTCTCCTCA |
| | | Reverse | **ACTTCTGTTCCATGTCGAC**CTAATCGTTAACAAGATGAGGAA |
| CrSBE | pESC-HIS | Forward | **GAGAAAAAACCCCGGATCC**ATGGATGAGATGATGAACTTCTC |
| | | Reverse | **ACTTCTGTTCCATGTCGAC**TTAATTTCCTTCAACTGCTGATGA |
| CrRS | pESC-HIS | Forward | **GAGAAAAAACCCCGGATCC**ATGGAAACTATCAACTTCTTCTC |
| | | Reverse | **ACTTCTGTTCCATGTCGAC**TCAATCTTGTGCAACAACCG |
| **Gene** | **Plasmid** | **Primer direction** | **Sequence (5‘-3‘)** |
| pTRV2 constructs for VIGS | | | |
| CrGO | pTRV2-MgChl | Forward | **ATATTGCTGCGGATC**CCTTTGATGCACGTGAAAATGGG |
| | | Reverse | **ATGCCCGGGCCTCGAG**GTCCAACGTTAGACTTAATTGACT |
| CrSBE | pTRV2-MgChl | Forward | **ATATTGCTGCGGATCC**GGCAAAAAATTACCACCGGG |
| | | Reverse | **ATGCCCGGGCCTCGAG**CAATTGCTTTAAGTAGTCACCA |
| **Gene** | **Plasmid** | **Primer direction** | **Sequence (5‘-3‘)** |
| Primers designed for mutagenesis (3Ω1 *Nicotiana benthamiana* transient expression) | | | |
| CrRS to GO CrRS-M1 (IC118-119MT, P220Δ, F375V) | 3Ω1 CrRS template | Forward | AAATTATGACTTACAATAATTTAGGCACAACT |
| | | Reverse | TATTGTAAGTCATAATTTCTGCCACACTA |
| | | Forward | TCAGCAGCAATCAGGTTTGAAGATCTTTTTCCA |
| | | Reverse | CCTGATTGCTGCTGATAATTTCGACACT |
| | | Forward | CTTCTAGTCCCAAGAGAAGC |
| | | Reverse | GCTTCTCTTGGGACTAGAAG |
| CrRS to GO CrRS-M2 (IC118-119MT, VSKLSAA213-219TSLLSGT, F375V) | 3Ω1 CrRS template | Forward | AAATTATGACTTACAATAATTTAGGCACAACT |
| | | Reverse | TATTGTAAGTCATAATTTCTGCCACACTA |
| | | Forward | ACGTCGTTACTGTCTGGAACACCAATCAGGTTTGAAGATCTT |
| | | Reverse | TTCCAGACAGTAACGACGTTTTTTTACACACTTTAATTAATTTTTCCCTT |
| | | Forward | CTTCTAGTCCCAAGAGAAGC |
| | | Reverse | GCTTCTCTTGGGACTAGAAG |
| CrRS to GO CrRS-M3 | 3Ω1 CrRS | Forward | AAATTATGTTATACAATAATTTAGGCACAACT |
| | | Reverse | CTAAATTATTGTATAACATAATTTCTGCCACACTA |
| | | Forward | AGCTCATCAGCCGTAGTAGAATGGG |

| | | | |
|---|---|---|---|
| (IC118-119MT, V311S, D313A) | template | Reverse | TACGGCTGATGAGCTTAGAGTTGCAGC |
| CrRS to GO CrRS-M4 (IC118-119MT, V311S, D313A) | 3Ω1 CrRS template | Forward | AAATTATGTTATACAATAATTTAGGCACAACT |
| | | Reverse | CTAAATTATTGTATAACATAATTTCTGCCACACTA |
| | | Forward | AGCTCATCAGCCGTAGTAGAATGGG |
| | | Reverse | TACGGCTGATGAGCTTAGAGTTGCAGC |
| | | Forward | ACCCACCTGCCCCTCTTC |
| | | Reverse | GAAGAGGGGCAGGTGGGT |
| CrGO to RS CrGO-M1 (V372F) | 3Ω1 CrGO template | Forward | CCTCTTTTATTCCCTAGAGAATGC |
| | | Reverse | GCATTCTCTAGGGAATAAAAGAGG |
| CrGO to SBE CrGO-M2 (A367P, V372F) | 3Ω1 CrGO template | Forward | CACCTGCTCCTCTTTTATTCCCT |
| | | Reverse | AGGGAATAAAAGAGGAGCAGGTG |
| CrSBE to RS CrSBE-M1 (ML113-114IC) | 3Ω1 CrSBE template | Forward | GCTAAGATCATCTGCTACAATAATTTG |
| | | Reverse | CAAATTATTGTAGCAGATGATCTTAGC |
| CrSBE to RS CrSBE-M2 (P372A) | 3Ω1 CrSBE template | Forward | TACACCCTGCTGCTCCC |
| | | Reverse | GGGAGCAGCAGGGTGT |
| **Gene** | **Plasmid** | **Primer direction** | **Sequence (5‘-3‘)** |
| Primers used for sequencing plasmid constructs | | | |
| | 3Ω1 | Forward | GATGAAAAAGCCCTAAAATTGGAG |
| | | Reverse | ATTATTCACAAATGAGAAACAGAATGG |
| | pESC-HIS | Forward | ATGATTTTTGATCTATTAACAGATA |
| | | Reverse | GTATAATGTTACATGCGTACAC |
| | pTRV2-MgChl | Forward | GATGGACATTGTTACTCAAGGAAGC |
| | | Reverse | CAGTCGAGAATGTCAATCTCGTAGG |

**Table S3.** VIGS fragment (300 bp) used for silencing.

| Gene | Nucleotide sequence |
|---|---|
| CrGO | CCTTTGATGCACGTGAAAATGGGCGAACGTTCGGCAATTATAGTATCAGATGCAAGAATG GCAAAAATAGTTCTTCATAATAACGGTTTAGCCGTTGCAGATCGGTCAGTAAATACTGTCG CAAGTATTATGACTTATAATAGTTTGGGTGTTACCTTTGCTCAGTATGGAGATTACTTAACA AAATTACGTCAAATCTATACTTTGGAACTTTTAAGTCAGAAAAAAGTTCGATCTTTCTACAG TTGTTTTGAAGATGAACTCGATACTTTTGTTAAGTCAATTAAGTCTAACGTTGGAC |
| CrSBE | GGCAAAAAAATTACCACCGGGTCCCAAAAAACTTCCTATTATAGGAAACCTCCATCAAATGA TAGGTTCACATCCTCATCGTGTTCTAAAAAAATTGGCCGATAAATATGGACCTATTATGCA CTTGCAAATGGGCCAACGATCAGCCGTTGTTATATCATCAGCGGAAAAAGCAAAAGAAAT ATTGAACATTTATGGTGTTCAAGTTGCTGATAGACCTCAACTCATCGTATCTAAAATTATGT TGTACAATAGTTTGGGTGTCACTTTTGCTCCTTATGGTGACTACTTAAAGCAATTGC |

**Table S4.** Primers used for qPCR analysis.

| Target | Primer direction | Sequence (5‘-3‘) |
|---|---|---|
| CrGO | Forward | TGGCCCGAACCAGAGAAGTT |
| | Reverse | AGCAGCCAACATAAACTCAGCA |
| CrSBE | Forward | CACCCTCCTGCTCCCATATTGT |
| | Reverse | ACTGAACCTCTCCGGCTCAA |
| q-Rps9 (reference gene) | Forward | TTGAGCCGTATCAGAAATGC |
| | Reverse | CCCTCATCAAGCAGACCATA |

Reference gene was used as described by Payne[22] and Li[22].

**Supplementary Figures**

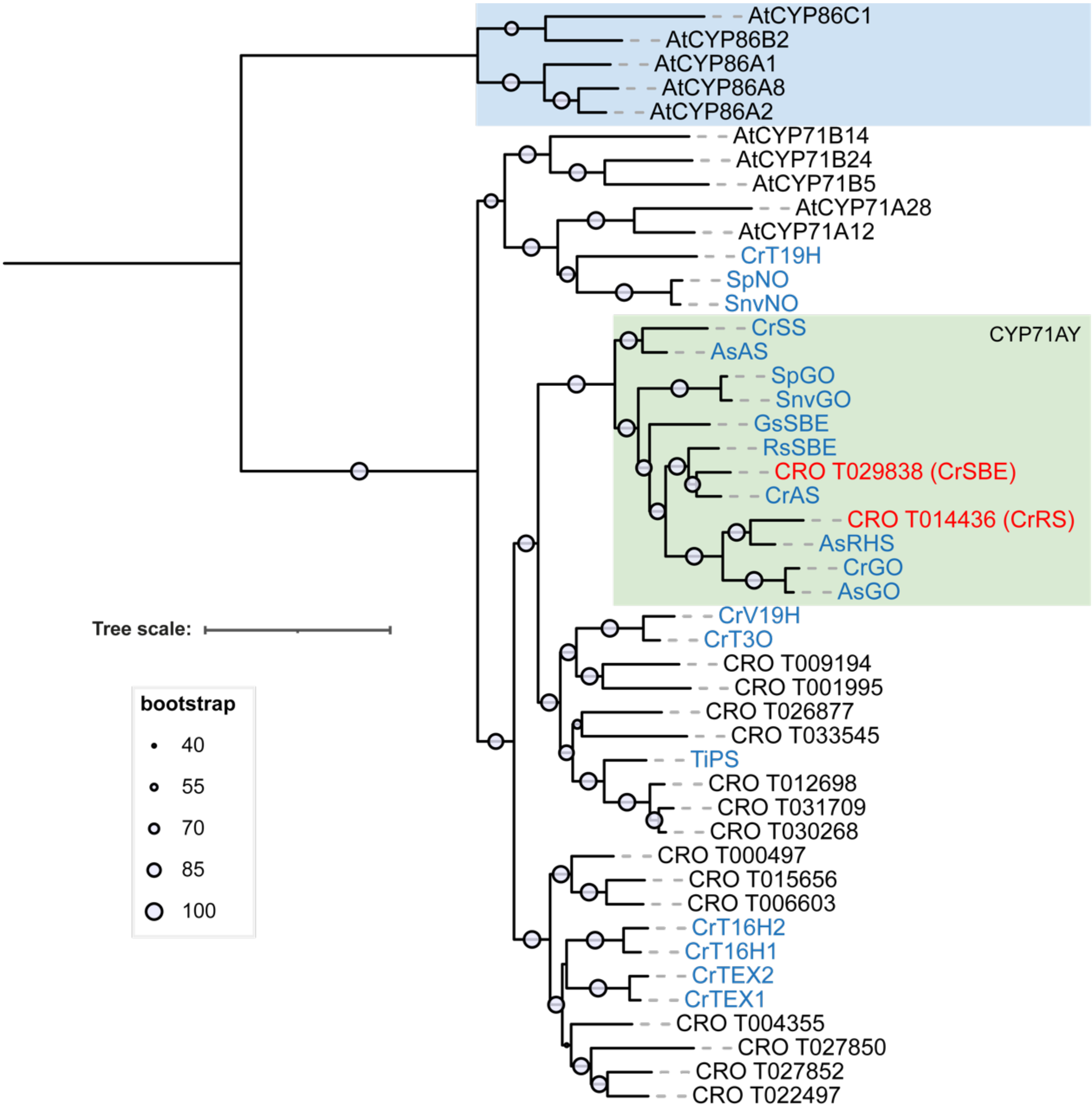


**Figure S1*. *Phylogenetic tree of *Catharanthus roseus* (CRO) cytochrome P450s from the CYP71 family.** Phylogenetic relationship of the cytochrome P450 (P450) enzymes identified from the CYP71 clan with previously characterized enzymes involved in MIA biosynthesis, all in the CYP71AY subfamily. The maximum-likelihood phylogenetic tree was constructed using iQtree[16] with default parameters and 1000 bootstraps (shown out of 100). The CYP71AY family investigated in this work is highlighted in green. P450s functionally characterized and shown to be involved in monoterpene indole alkaloid biosynthesis are coloured in blue. *Arabidopsis thaliana* (At) CYP86 clan 86 was used to root the tree. The phylogenetic tree was visualized and inferred using iToL[17].

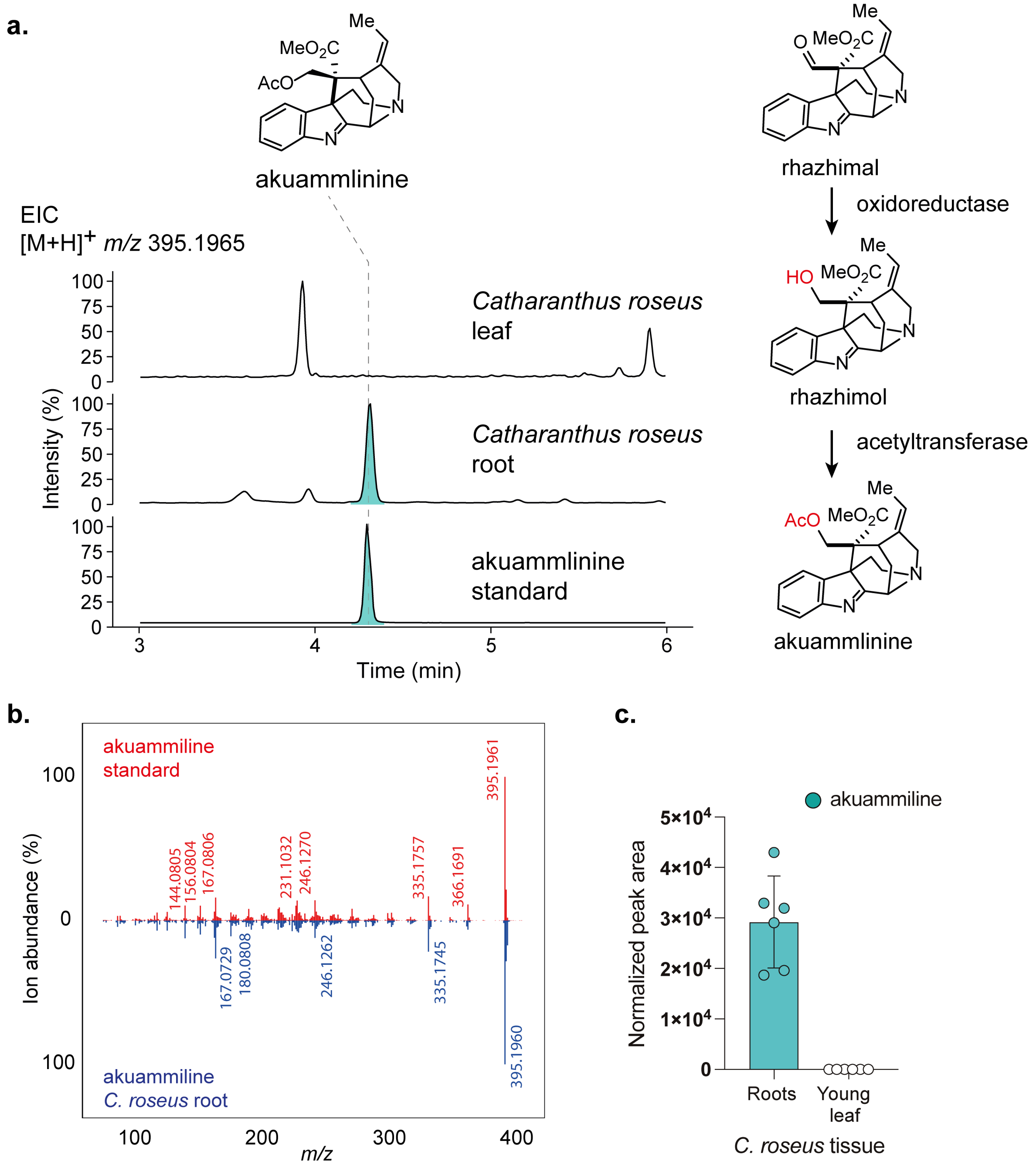


**Figure S2. Detection of akuammiline in *C. roseus* tissues.** (**a**) LC-MS profiles of *C. roseus* root and young leaf tissue showing accumulation of akuammiline. Akuammiline is present in the roots of *C. roseus* and is not detected in the young leaves. Akuammiline biosynthesis starts with rhazimal, where an oxidoreductase reduces the aldehyde to form rhazhimol, and subsequent acetylation leads to the formation of akuammiline. While akuammiline biosynthesis has been elucidated in the plant *Alstonia scholaris*[8], this compound was not previously noted to be present in *C. roseus*. (**b**) $MS^2$ spectra of akuammiline authentic standard (red) compared to analyte detected in *C. roseus* roots (blue) show identical spectral match. (**c**) Normalized peak area of akuammiline detected in the roots of 6 independent *C. roseus* plants.

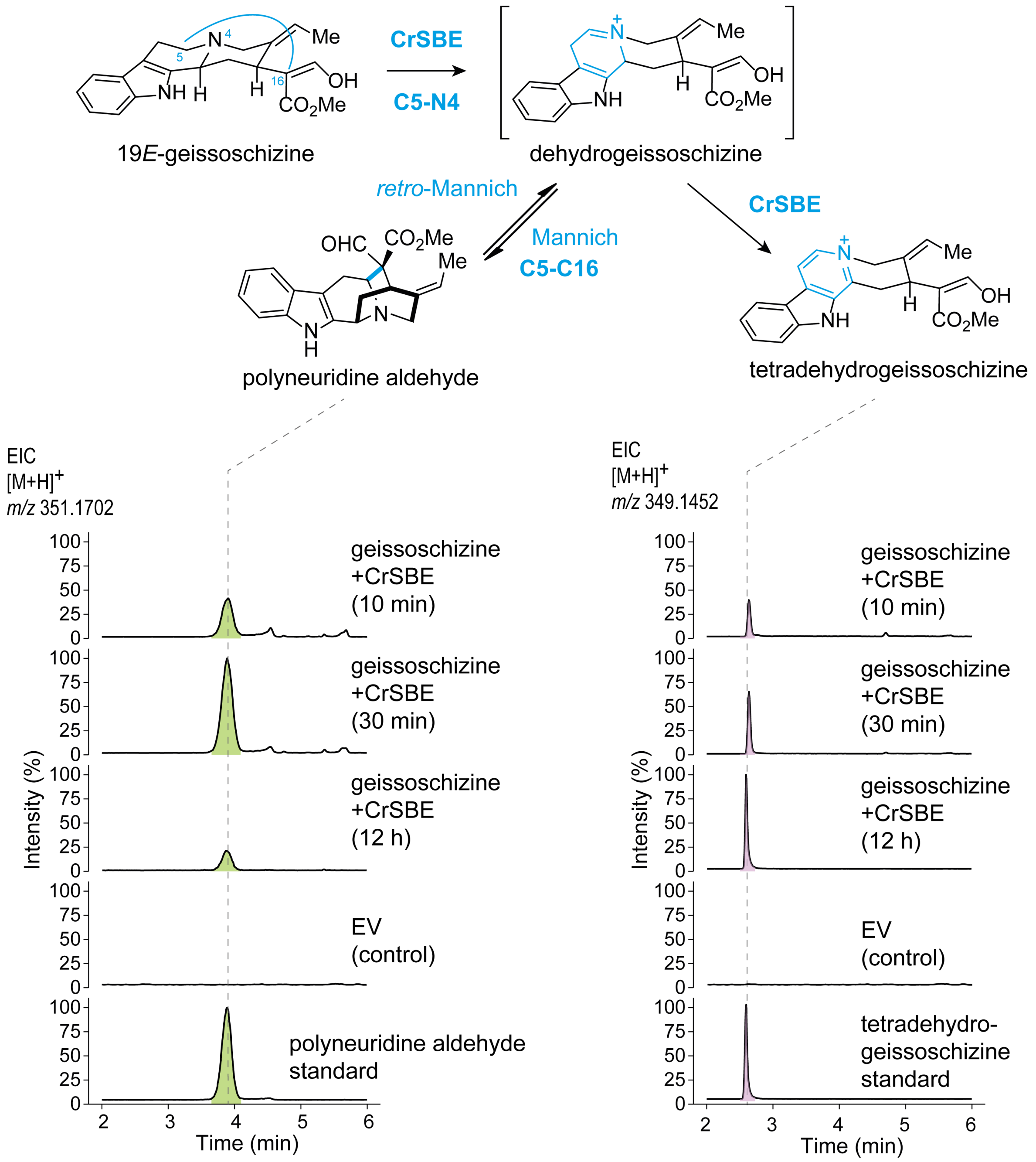


**Figure S3. In vitro reactions of CrSBE (CRO_T029838).** In vitro assays using yeast microsomal preparations harbouring CrSBE. These microsomes were incubated with 19*E*-geissoschizine and aliquots were quenched at 10 min, 30 min, and 12 h. The dehydrogeissoschizine iminium intermediate appears to be in an equilibrium with polyneuridine aldehyde via a Mannich-type rearrangement over the short-time course experiment. In prolonged incubation only tetra-dehydro-geissoschizine is observed.

**a.**

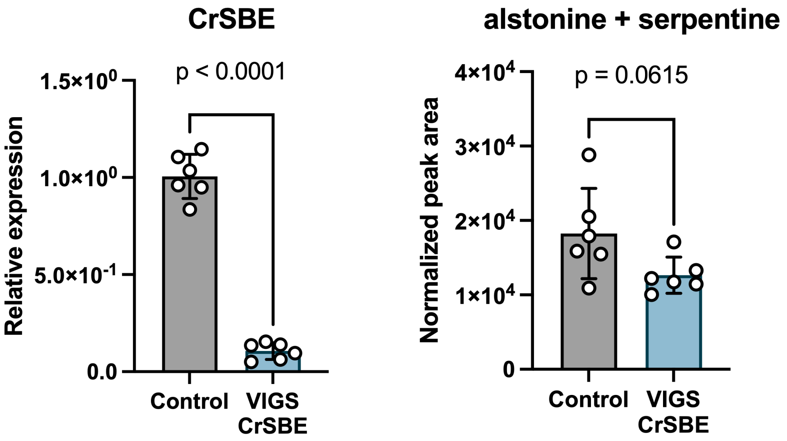


**b.**

| amino acid sequence similarity | CRO_T029838 (CrSBE) | CrSS | CrAS |
|---|---|---|---|
| CRO_T029838 (CrSBE) | | 49.6 | 78.9 |
| CrSS | 49.6 | | 50.8 |
| CrAS | 78.9 | 50.8 | |

**c.**

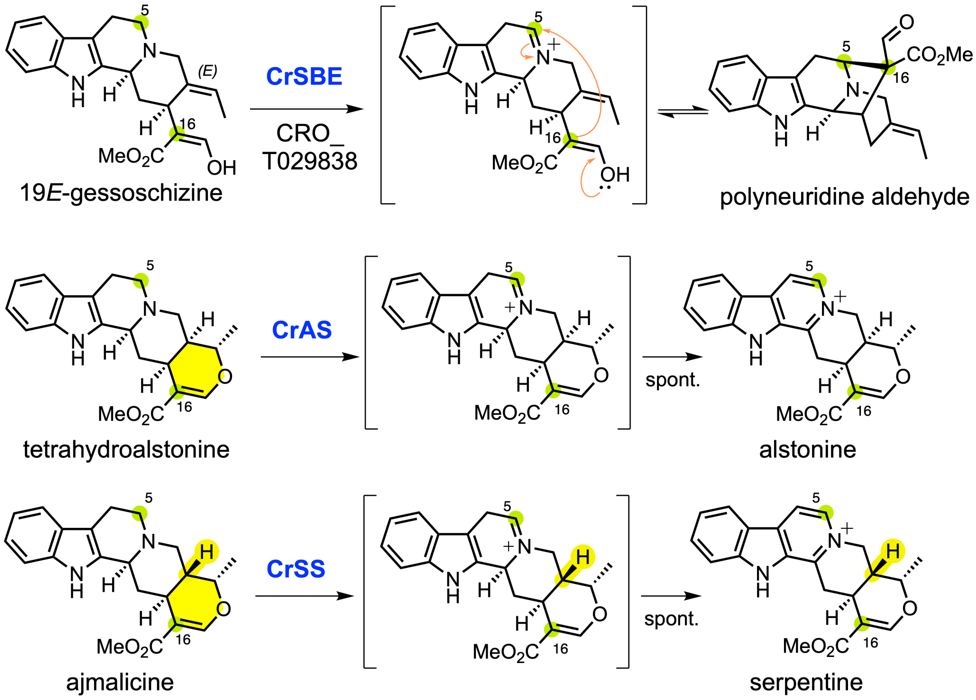


**Figure S4. VIGS of CrSBE (CRO_T029838) in *C. roseus*.** **(a)** Effective silencing of the CrSBE gene in *C. roseus* va. SA leaves. Despite statistically significant silencing of CrSBE, no statistically significant changes in alstonine or serpentine metabolite levels were observed. The values represent mean ± standard deviation (SD) (n = 6), two-tailed Student's *t*-test. (**b**) Amino acid similarity of the CrSBE (CRO_T029838) against previously characterised *C. roseus* SBE enzymes, AS (alstonine synthase), and SS (serpentine synthase). (**c**) Reactions catalyzed by the CrSBE (CRO_T029838) reported in this study and CrAS. The substrates of CrSBE and CrAS/CrSS share structural similarities, but the aromatization of the E ring (yellow) generates different oxidative rearrangements.

| | CRO_T014436 (CrRS) | CRO_T029838 (CrSBE) | AsRHS | GsSBE | RsSBE | CrAS | AsAS | CrGO | SnvGO | AsGO | SpGO | CrSS |
|---|---|---|---|---|---|---|---|---|---|---|---|---|
| CRO_T014436 (CrRS) | | 51.3 | 73.5 | 50.5 | 51.6 | 51.2 | 48.9 | 58.3 | 47.7 | 59.3 | 49.1 | 44.0 |
| CRO_T029838 (CrSBE) | 51.3 | | 53.9 | 58.7 | 73.6 | 78.9 | 58.1 | 50.9 | 55.7 | 52.3 | 56.6 | 49.6 |
| AsRHS | 73.5 | 53.9 | | 53.1 | 55.3 | 55.0 | 49.4 | 61.6 | 49.5 | 62.4 | 50.4 | 44.9 |
| GsSBE | 50.5 | 58.7 | 53.1 | | 59.4 | 59.2 | 55.5 | 50.9 | 54.5 | 51.3 | 54.6 | 52.0 |
| RsSBE | 51.6 | 73.6 | 55.3 | 59.4 | | 74.1 | 58.0 | 52.0 | 53.7 | 52.6 | 55.5 | 51.2 |
| CrAS | 51.2 | 78.9 | 55.0 | 59.2 | 74.1 | | 58.7 | 51.5 | 53.6 | 52.6 | 54.7 | 50.8 |
| AsAS | 48.9 | 58.1 | 49.4 | 55.5 | 58.0 | 58.7 | | 47.7 | 54.9 | 48.8 | 55.8 | 67.8 |
| CrGO | 58.3 | 50.9 | 61.6 | 50.9 | 52.0 | 51.5 | 47.7 | | 46.8 | 90.2 | 47.8 | 44.6 |
| SnvGO | 47.7 | 55.7 | 49.5 | 54.5 | 53.7 | 53.6 | 54.9 | 46.8 | | 47.6 | 92.1 | 50.0 |
| AsGO | 59.3 | 52.3 | 62.4 | 51.3 | 52.6 | 52.6 | 48.8 | 90.2 | 47.6 | | 48.8 | 45.6 |
| SpGO | 49.1 | 56.6 | 50.4 | 54.6 | 55.5 | 54.7 | 55.8 | 47.8 | 92.1 | 48.8 | | 49.9 |
| CrSS | 44.0 | 49.6 | 44.9 | 52.0 | 51.2 | 50.8 | 67.8 | 44.6 | 50.0 | 45.6 | 49.9 | |

**Figure S5. Sequence similarity of CrRS and CrSBE to known, functionally characterized CYP71AY proteins.** The percentage of amino acid sequence similarity of CrRS (CRO_T014436) and CrSBE (CRO_T029838) to that of functionally characterized CYP71AY cytochrome P450 enzymes involved in monoterpene indole alkaloid biosynthesis.

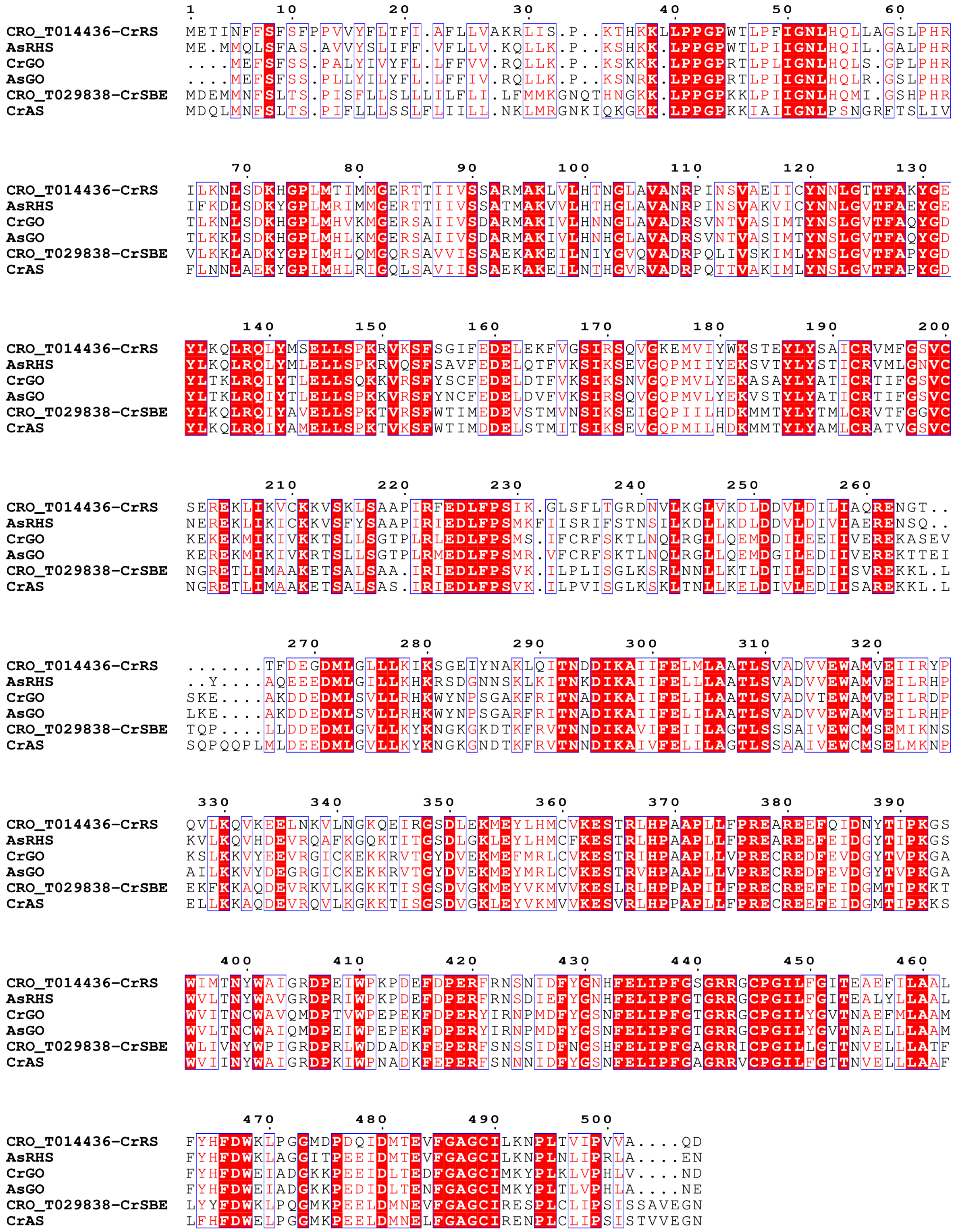


**Figure S6. Multiple sequence alignments of CrRS and CrSBE.** MUSCLE[14] amino acid alignments of CrGO, CrRS (CRO_T014436), CrSBE (CRO_T029838), *Alstonia scholaris* geissoschizine oxidase (AsGO), *Alstonia scholaris* rhazimal synthase (AsRHS), and *Catharanthus roseus* Alstonine synthase (CrAS). Sequences in Figure S5 with the highest amino acid identity to CrGO, CrRS and CrSBE were selected for alignment. Alignment was visualised using ESPript[30] webserver.

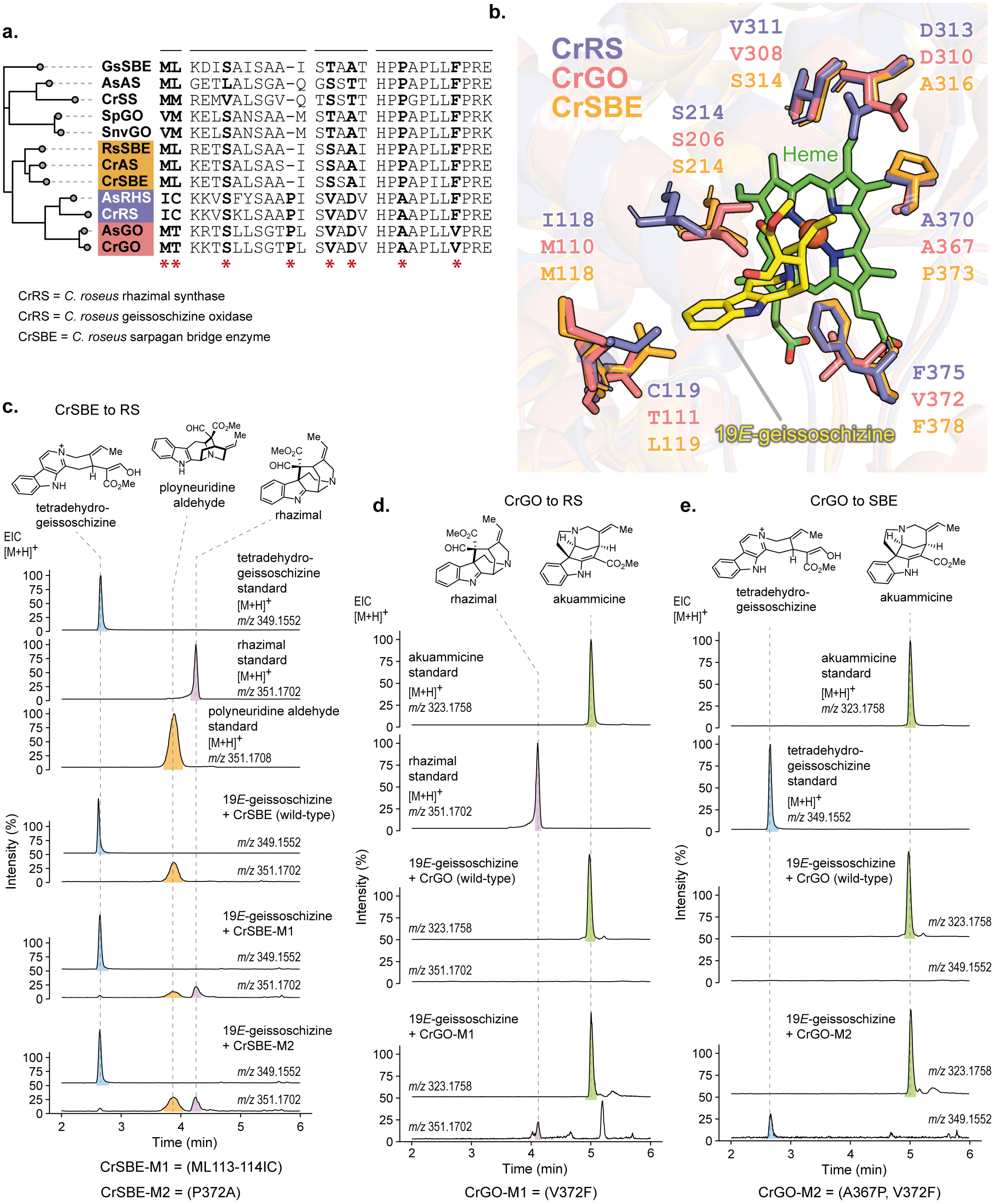


**Figure S7. Site directed mutagenesis of CrSBE and CrGO alters enzymatic activity on the MIA pathway.** (**a**) Phylogenetic relationship of functionally characterized GO, SBE and SBE enzymes. (**b**) Superimposed AlphaFold3 models of CrRS (purple), CrGO (pink), and CrSBE (orange) catalytic pocket reveals key residue interactions with the 19*E*-geissoschizine (yellow sticks) substrate. The catalytic heme is presented in green. (**c**) EICs show enzymatic activity of CrSBE mutants, M1 and M2 on 19*E*-geissoschizine. Wild-type CrSBE converts 19*E*-geissoschizine to tetradehydrogeissoschizine, whereas CrSBE-M1 and CrSBE-M2 show altered activity, with CrSBE-M2 producing rhazimal and polyneuridine aldehyde. (**d**) EICs of CrGO mutant M1 on 19*E*-geissoschizine. CrGO-M1 produces low amounts of rhazimal along with akuammicine. (**e**) EICs of CrGO mutant M2 on 19*E*-geissoschizine. CrGO-M2 is able to produce low levels of tetradehydrogeissoschizine along with akuammicine. Enzymatic activity presented (**c**), (**d**), and (**e**) of the wild-type and mutant proteins were performed in *N. benthamiana*. Abbreviations; EIC, Extracted ion chromatograms.

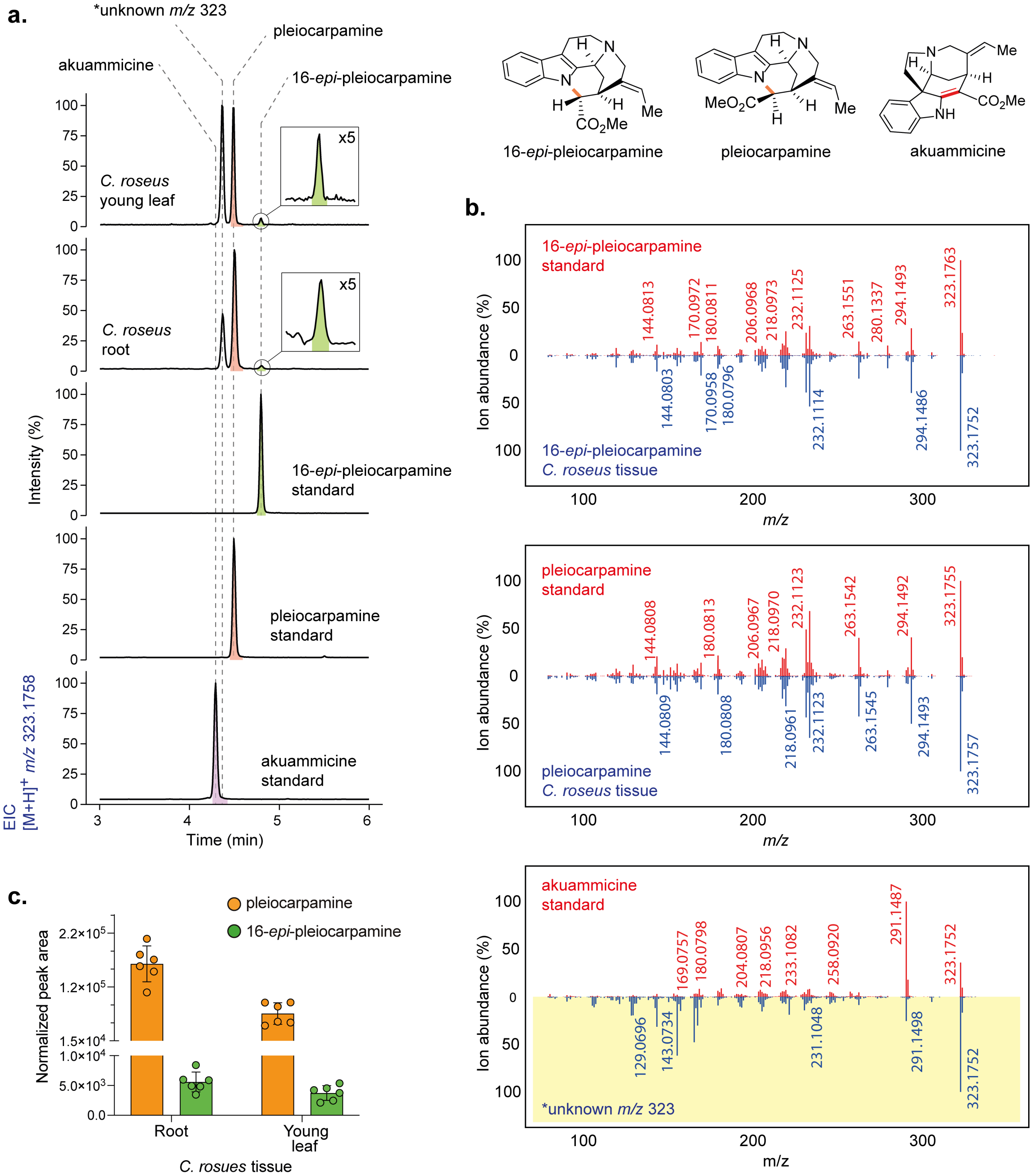


**Figure S8. Metabolite analysis of pleiocarpamine and 16-*epi*-pleiocarpamine in *C. roseus* tissues.** (**a**) LC-MS profiles of *C. roseus* root and young leaf tissue showing the accumulation of pleiocarpamine and 16-*epi*-pleiocarpamine. Pleiocarpamine is present at higher levels than 16-*epi*-pleiocarpamine. An unknown compound (*m/z* 323) was recorded with elution times similar to akuammicine. (**b**) MS$^2$ spectra of pleiocarpamine, 16-*epi*-pleiocarpamine, and akuammicine authentic standards (red) compared to compounds detected in *C. roseus* tissue (blue). Pleiocarpamine and 16-*epi*-pleiocarpamine showed identical spectral match. The unknown compound (*m/z* 323) observed in the young leaf did not show MS$^2$ spectra similarity to the any of the authentic standards. (**c**) Normalized peak areas of pleiocarpamine and 16-*epi*-pleiocarpamine detected in the roots and young leaves of 6 independent *C. roseus* plants. Peak areas were normalized to internal standard tabersonine. Error bars represent ± standard deviation (SD) of the mean.

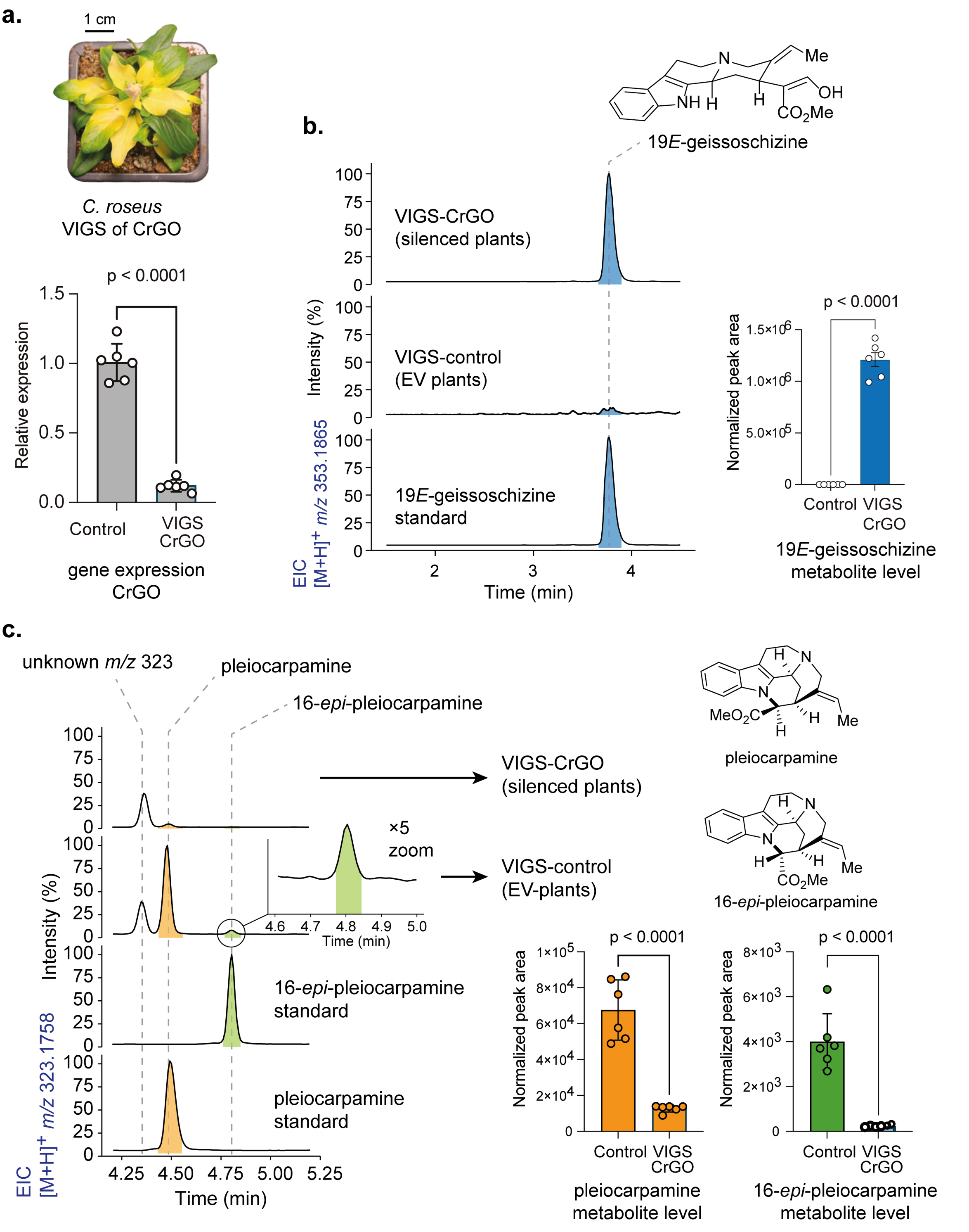


**Figure S9. VIGS of CrGO in *C. roseus*.** (**a**) The gene encoding CrGO exhibits a much lower expression value compared to the empty vector (EV) control, indicating successful silencing. (**b**) Silencing GO in the leaves led to the accumulation of the canonical GO substrate, 19*E*-geissoschizine, which further demonstrate the effective silencing of the gene. Representative LC-MS extracted ion chromatogram (EIC) is shown alongside the bar graph for metabolite accumulation. (**c**) Metabolite levels of the GO knockdown tissue in comparison to controls. Representative LC-MS extracted ion chromatogram (EIC) is shown alongside the bar graph for metabolite accumulation The values represent mean ± standard deviation (SD) (n = 6), two-tailed Student's *t*-test.

### Supporting NMR Data

### NMR analysis of strictamine

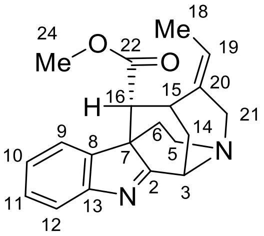

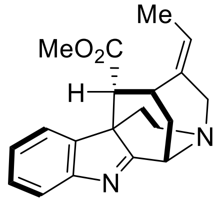

$^{1}$H-$^{1}$H COSY
bold line

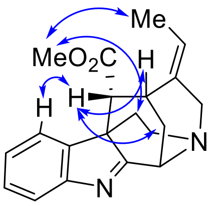

NEOSY

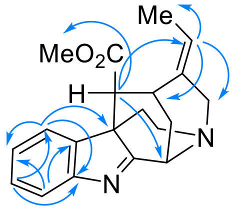

HMBC
(H to C)

$^{1}$H NMR (700 MHz, $CDCl_3$) δ 7.71 (d, *J* = 7.8 Hz, 1H, H-12), 7.47 – 7.41 (m, 1H, H-9), 7.41 (d, *J* = 7.5 Hz, 1H, H-11), 7.29 – 7.24 (m, 1H H-10), 5.84 (d, *J* = 7.5 Hz, 1H, H-19), 5.29 (s, 1H, H-3), 4.43 (d, *J* = 15.2 Hz, 1H, H-21), 3.75 (s, 3H, H-24), 3.78-3.73 (m, 1H, H-6), 3.68 (s, 1H, H-15), 3.65 (d, *J* = 13.8 Hz, 1H, H-21), 3.26 – 3.19 (m, 1H, H-5), 2.90 (d, *J* = 15.6 Hz, 1H, H-5), 2.82 (d, *J* = 14.4 Hz, 1H, H-14), 2.20 (dd, *J* = 15.8, 5.1 Hz, 1H, H-6), 2.15 (d, *J* = 3.4 Hz, 1H, H-16), 2.01 (d, *J* = 13.3 Hz, 1H, H-14), 1.65 (d, *J* = 6.6 Hz, 3H, H-18).

$^{13}$C NMR (176 MHz, $CDCl_3$) δ 192.3 (C-2), 170.5 (C-22), 154.7 (C-13), 144.3 (C-8), 129.2 (C-11), 127.3 (C-19), 127.2 (C-10), 127.0 (C-20), 123.5 (C-9), 122.4 (C-12), 55.1 (C-3), 54.9 (C-7), 54.1 (C-16), 52.7 (C-21), 52.1 (C-24), 51.3 (C-5), 33.7 (C-14), 31.0 (C-15), 25.5 (C-6), 13.4 (C-18).

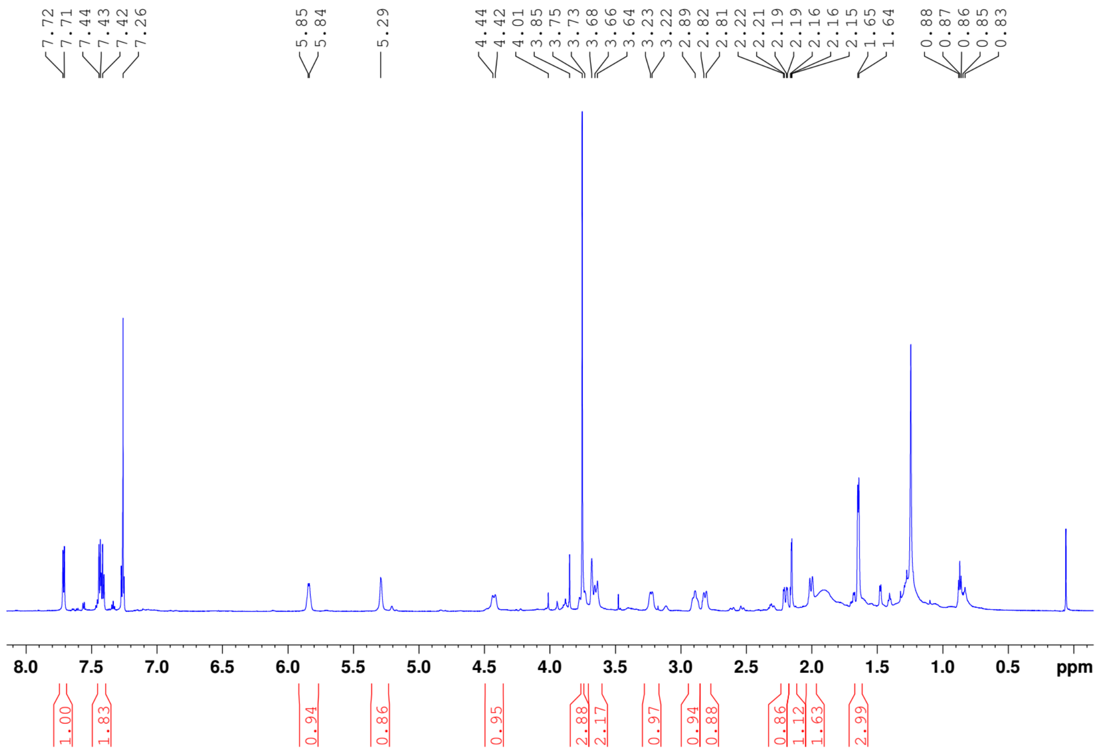


**NMR Figure S1.** $^{1}$H-NMR of strictamine.

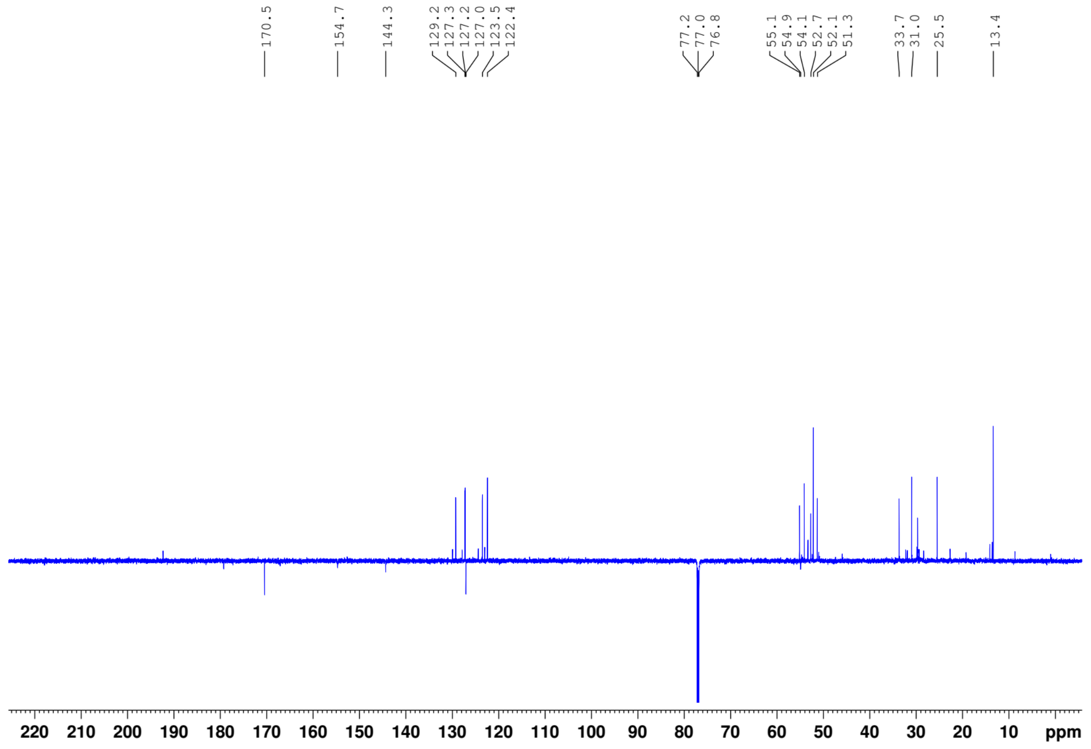


**NMR Figure S2.** $^{13}$C-NMR of strictamine

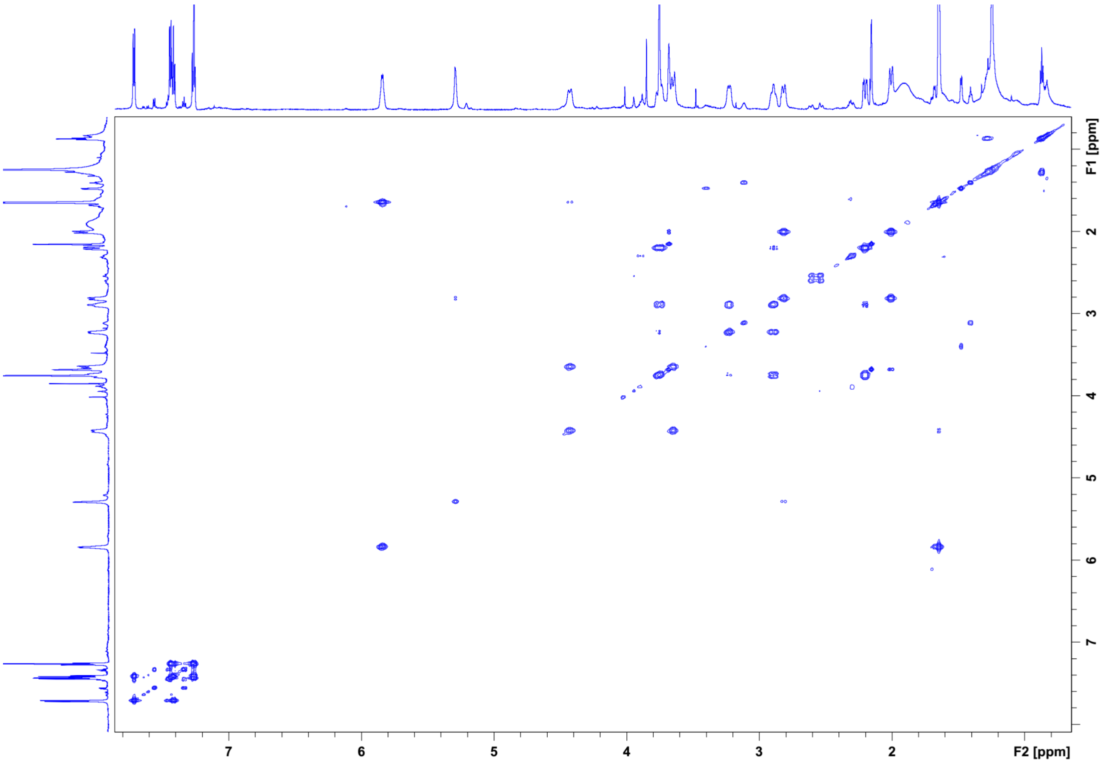


**NMR Figure S3.** $^{1}H$-$^{1}H$ COSY of strictamine.

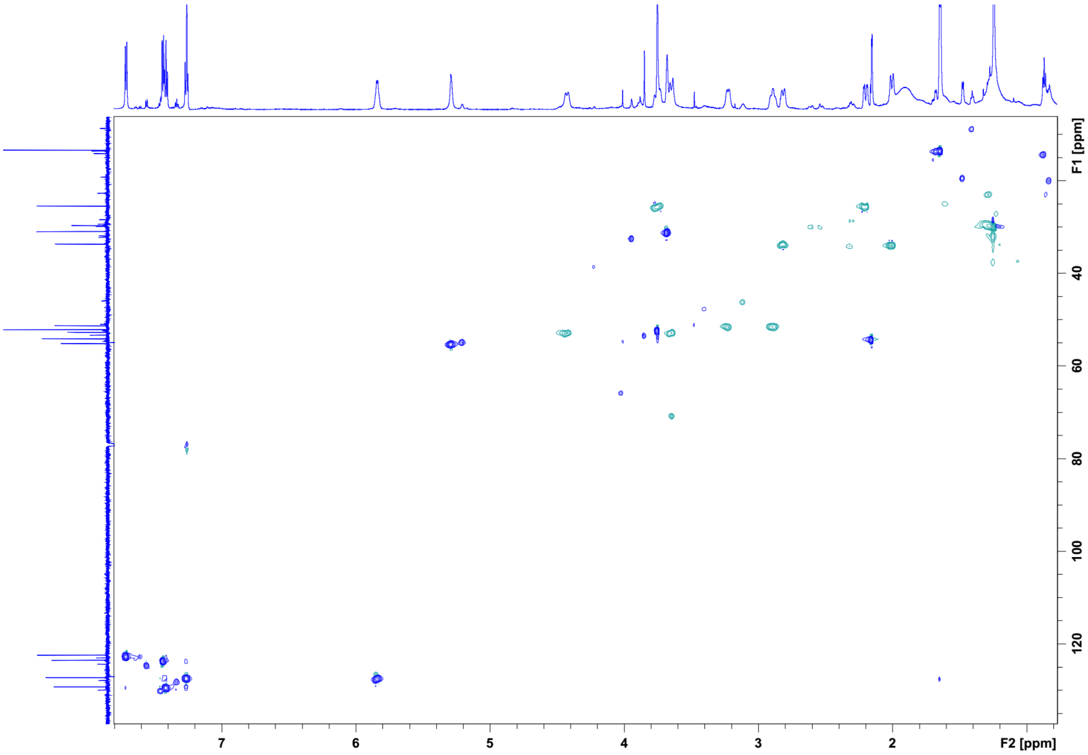


**NMR Figure S4.** HSQC of strictamine.

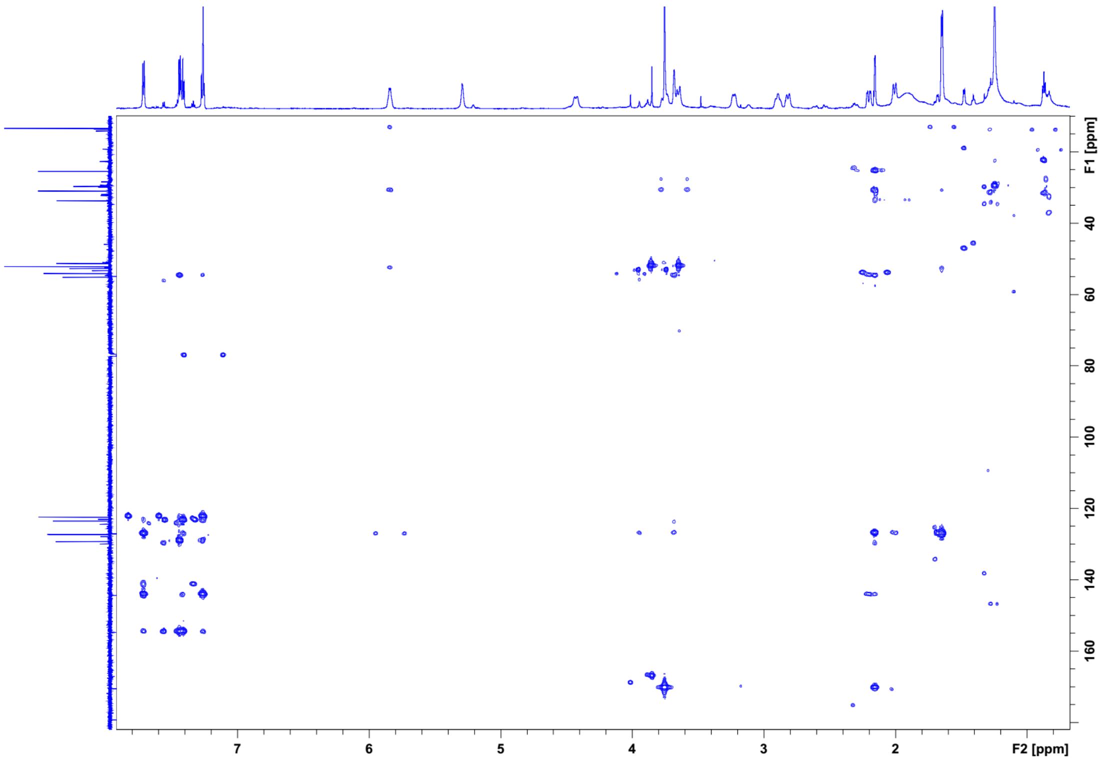


**NMR Figure S5.** HMBC of strictamine.

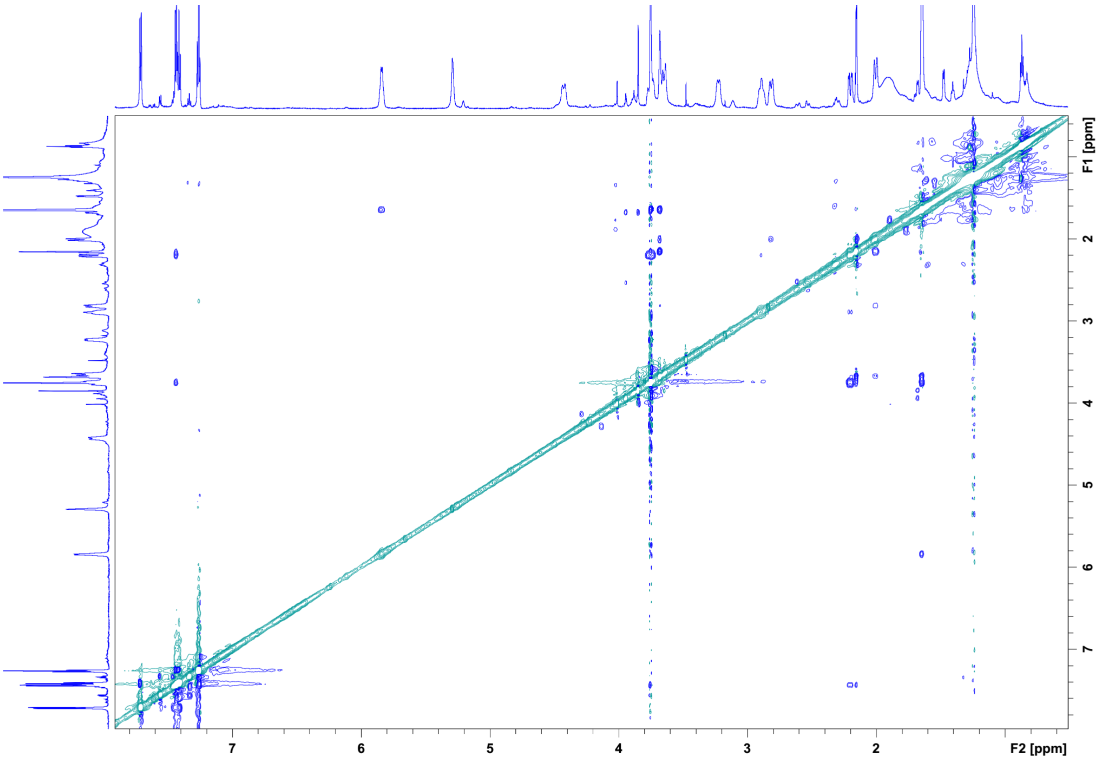


**NMR Figure S6.** ROESY of strictamine.

#### Partial NMR analysis of rhazimal

$^1$H-NMR (700 MHz, $CDCl_3$) δ 8.49 (s, 1H), 7.72 (d, *J* = 7.7 Hz, 1H), 7.56 (d, *J* = 7.4 Hz, 1H), 7.46 (t, *J* = 7.7 Hz, 1H), 7.34 (t, *J* = 7.4 Hz, 1H), 5.87 (br, s, 1H), 5.22 (br, s, 1H), 4.48 (br, s, 1H), 4.00 – 3.87 (m, 2H), 3.85 (s, 3H), 2.91 (br, s, 1H), 2.71 – 2.45 (m, 3H), 2.39 – 2.21 (m, 2H), 1.70 (d, *J* = 15.8 Hz, 3H).

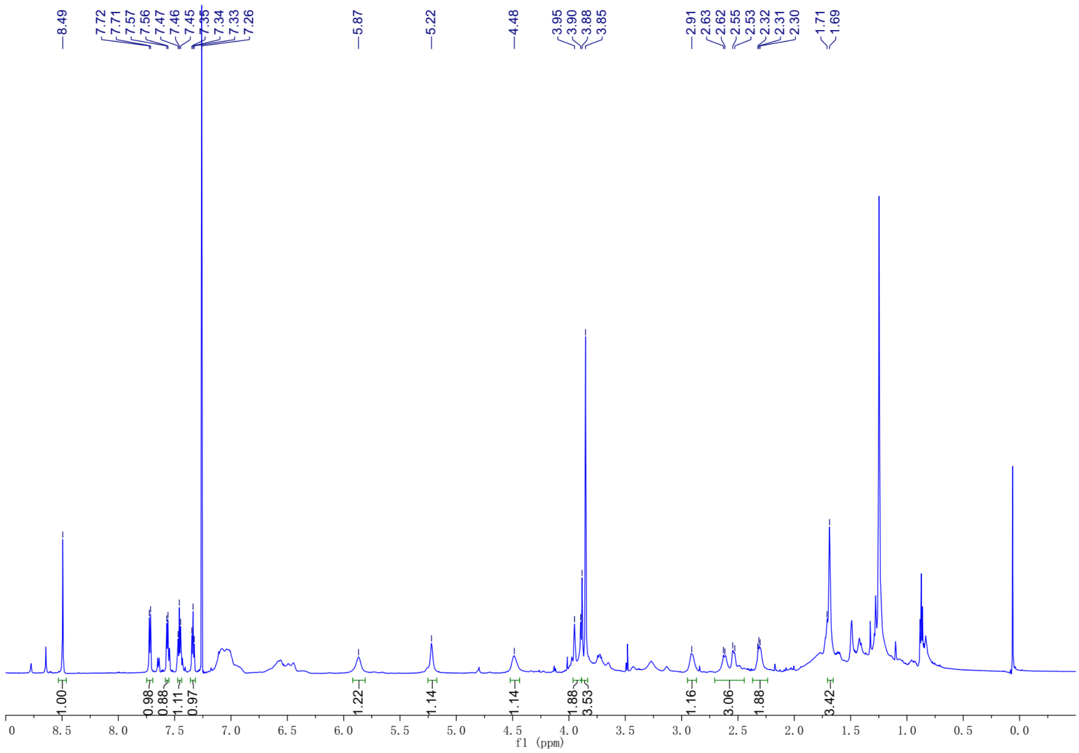


**NMR Figure S7.** $^1$H-NMR of rhazimal

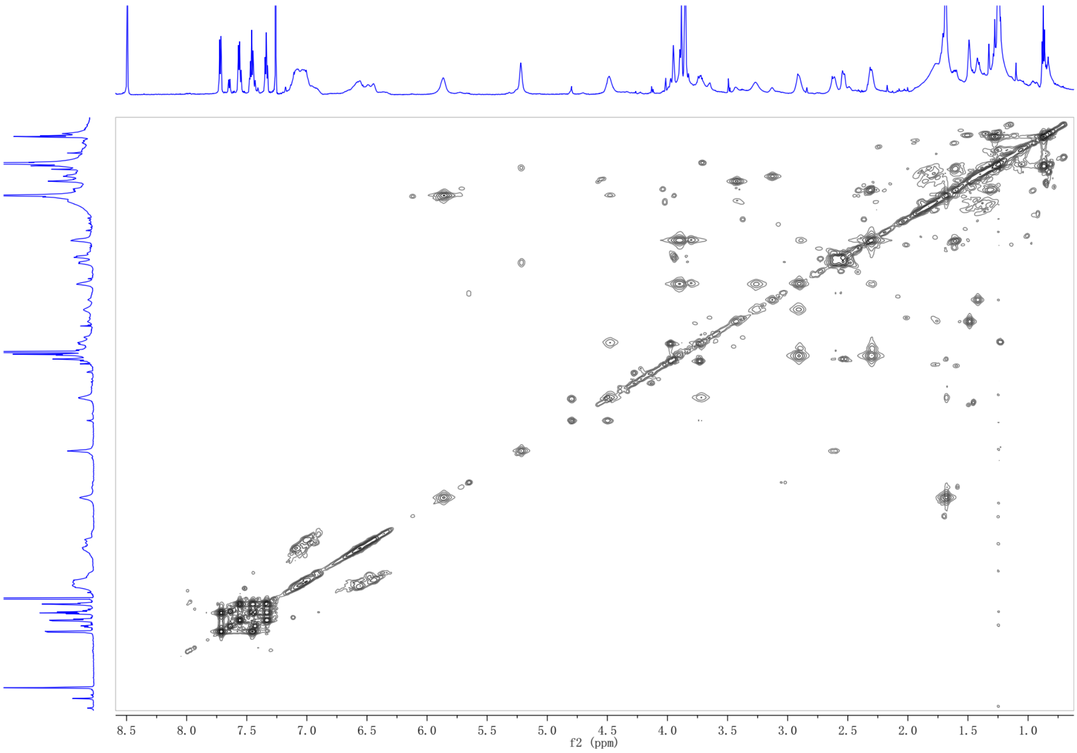


**NMR Figure S8.** $^{1}H$-$^{1}H$ COSY of rhazimal

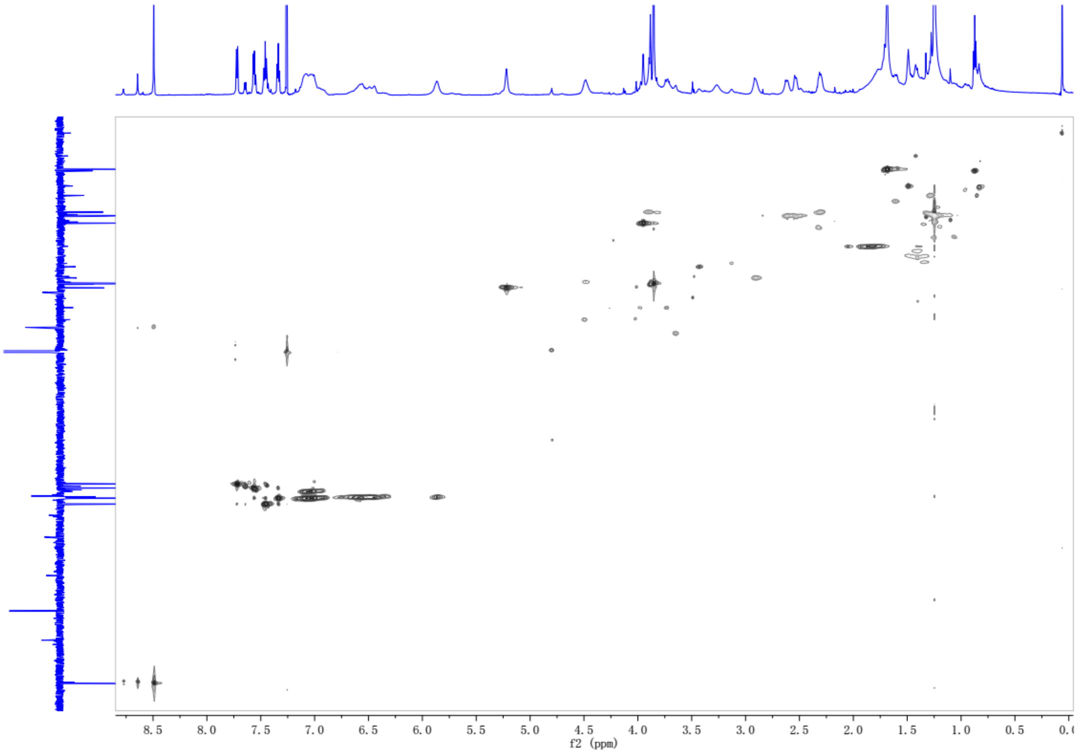


**NMR Figure S9.** HSQC of rhazimal
